# Sculpting rupture-free nuclear shapes in fibrous environments

**DOI:** 10.1101/2021.10.19.465049

**Authors:** Aniket Jana, Avery Tran, Amritpal Gill, Rakesh K. Kapania, Konstantinos Konstantopoulos, Amrinder S. Nain

**Author notes:** Corresponding author: Amrinder S. Nain, **Email:**. equal second authors (Avery Tran, Amritpal Gill).

## Abstract

Cytoskeleton-mediated force transmission regulates nucleus morphology. How nuclei shaping occurs in fibrous in vivo environments remains poorly understood. Here a suspended nanofiber assay of precisely-tunable (nm-μm) diameters is used to quantify nucleus plasticity in fibrous environments mimicking the natural extracellular matrix. In contrast to the apical cap over the nucleus in cells on 2-dimensional surfaces, the cellular cytoskeleton of cells on fibers displays a uniform actin network caging the nucleus. The role of contractility-driven caging in sculpting nuclear shapes is investigated as cells spread on aligned single fibers, doublets, and multiple fibers of varying diameters. Cell contractility increases with fiber diameter due to increased focal adhesion clustering and density of actin stress fibers, which correlates with increased mechanosensitive transcription factor YAP translocation to the nucleus. Unexpectedly, large- and small-diameter fiber combinations lead to teardrop-shaped nuclei due to stress-fiber anisotropy across the cell. As cells spread on fibers, diameter-dependent invaginations that run the nucleus’s length are formed at contact sites. The deepest and sharpest invaginations are insufficient to trigger nucleus rupture, often observed in 2D or confined systems. Overall, we describe the unknown adaptability of nuclei to fibrous environments and resultant sculpting of the nucleus shapes, with pathophysiological implications.

## 1. Introduction

The nucleus, as the stiffest cellular organelle, plays a central role in mechanotransduction ^[1,2]^. Proper control of nuclear shapes is one of the primary cellular functions controlling cell migration, differentiation, and tissue morphogenesis ^[3]^. Thus, not surprisingly, defects in nuclear shapes are often implicated in various disease states, including progeria, muscular dystrophy, and cancer metastasis ^[4]^. Mechanical forces regulate nuclear shapes ^[5–7]^ originating from cells’ dynamic interactions with their surrounding microenvironment, i.e., the extracellular matrix (ECM), and transmitted to the nucleus by the cell cytoskeleton. The nuclear envelope composed of the lamin intermediate filament networks is critical in sustaining external forces exerted on the nucleus. Lamin network mutations and deficiencies cause significant reduction of the nuclear stiffness ^[8]^, leading to nuclear blebbing ^[9]^, and mislocalization of DNA repair factors, and DNA damage ^[10]^.

Mechanical forces primarily act through the actomyosin contractions in the stress fibers to control nucleus shape to induce nucleo-cytoplasmic shuttling of transcription factors like YAP/TAZ, MKL1, and chromatin remodeling proteins such as HDAC ^[11,12]^. The force-mediated nuclear shape changes also induce reorganization and stretching of the internal chromatin domains ^[13]^, resulting in altered transcriptional activity and mechanical properties of the nucleus ^[14]^. A growing consensus in the field has been that nuclei rupture as they undergo drastic shape changes, causing mislocalization of signaling molecules (6,12). Large scale mechanical forces, driven by ECM stiffness, mold cell shape, which controls the nucleus shape; high aspect ratio nuclei induce high curvature at the poles, making them susceptible to rupture ^[9]^.

*In vivo* 3D aligned matrices, frequently observed in healthy and diseased states, including tendons ^[16,17]^, muscle tissue^[18,19]^, and extracellular regions surrounding metastatic tumors ^[20–22]^, provide topographic cues for cells to spread uniaxially and migrate persistently by exerting forces ^[23,24]^. Fibrillar matrices composed of individual collagen fibrils range in size from 70 to 300 nm ^[25–28]^ that can bundle to larger fibers varying from 1 to 20 μm in diameter ^[26,28]^. Mimicking the native ECM architecture within an in vitro setting is often highly challenging, and flat 2D substrates with and without anisotropic features and 3D gels have been used extensively to study cell behavior^[29]^. 2D systems have limited physiological relevance. While 3D collagen gels capture the fibrillar architecture, the inherent heterogeneity in these matrices renders it difficult to understand the role of fiber dimensions and organization in regulating cytoskeletal and nuclear responses ^[30,31]^. Studies by the Yamada group have demonstrated how single-cell behavior in 3D microenvironments can be recapitulated through the use of narrow 1D microprinted lines of varying widths (1-40 μm) ^[32]^. Microprinted lines are essentially 2D surfaces, and in our study, we inquired if suspended 1D fibrillar architecture of precisely controlled fiber diameters (150-6000 nm) and interfiber spacing regulated nuclear responses of uniaxial spread cells, as we have previously shown protrusive, contractile, and migratory behavior of uniaxial cells to be sensitive to fiber curvature and spacing ^[33–39]^. In this study, we chose nucleus rupture and the spatial localization of YAP/TAZ as two markers of nuclear response to changes in fiber curvatures. We discovered that cytoskeletal and lamin networks in suspended cells are localized in an almost uniform caging structure surrounding the nucleus, contrary to the preferential apical localization in flat continuous surfaces. Fiber-curvature driven cytoskeleton tension led to precise sculpting of nucleus shape, including unique teardrop shapes due to actin stress fiber anisotropy and invaginations that ran the length of the nucleus. We found that all nucleus shapes, including the sharpest invaginations formed on nanofibers, did not undergo rupture events, indicating remarkable adaptability of nuclei to fibrillar environments. Nuclear translocation of YAP increased with the diameter or fiber density. Overall, we describe cellular mechanosensitivity unique to fiber matrices with implications in pathophysiology.

## 2. Results

### 2.1. Suspended cells show uniform nuclear caging of cytoskeletal elements and lamins

We wished to compare the responses of cells plated on flat 2D as opposed to those attached to suspended fibers of varying diameters. We utilized force-measuring nanonets composed of large diameter (~2000 nm) ‘strut’ fibers fused to orthogonal small diameter aligned fibers (Nanonet force Microscopy, NFM) ^[38,40–42]^. To investigate the role of fiber diameter, we selected three different fiber diameters: 200 nm, 350 nm, and 800 nm, while keeping the spacing between large strut fibers constant (~250 μm). Fiber beam stiffness, midspan of a beam scales approximately with ~d^2^; thus, constant beam length allows us to directly compare the effects of fiber diameter contributing to an increase in stiffness ^[33,38,40]^. We investigated the dynamics of cell spreading and the organization of cytoskeletal networks using a custom setup to add cell suspension droplets on the fibronectin-coated fiber networks (**Figure 1a, b**). The inter-fiber spacing (10-12 μm) was chosen to ensure that cells spread along two fibers to form symmetric parallel-cuboidal shapes (**Figure 1b (iii)**). We monitored cell spreading using optical microscopy for cells attached approximately midway across the fiber span length (**Figure 1c**). After making initial contact with the fiber networks, cells began to protrude along the nanofibers (**Movie S1**). Consistent with our previous findings^[34]^, we observed protrusions formed during spreading were primarily actin-based, while microtubules and vimentin intermediate filaments localized later during the spreading cycle (**Figure S1 b**). To characterize cell size and shape during spreading, we tracked the cell projected area and the circularity for one hour for the three diameters (**Movies S2-4**). The cell area growth curves for all the fiber diameter categories showed a faster initial incremental phase followed by gradual maturation at longer times (**Figure 1d (i, iii)**), matching growth rate kinetics on 2D substrates^[43]^. Interestingly, we found that cells achieved steady-state area and circularity faster (~ 40 min) on 200 nm diameter fibers, while cells on larger diameter fibers demonstrated higher spread areas and circularity (**Figure 1d**).

**Figure 1:**
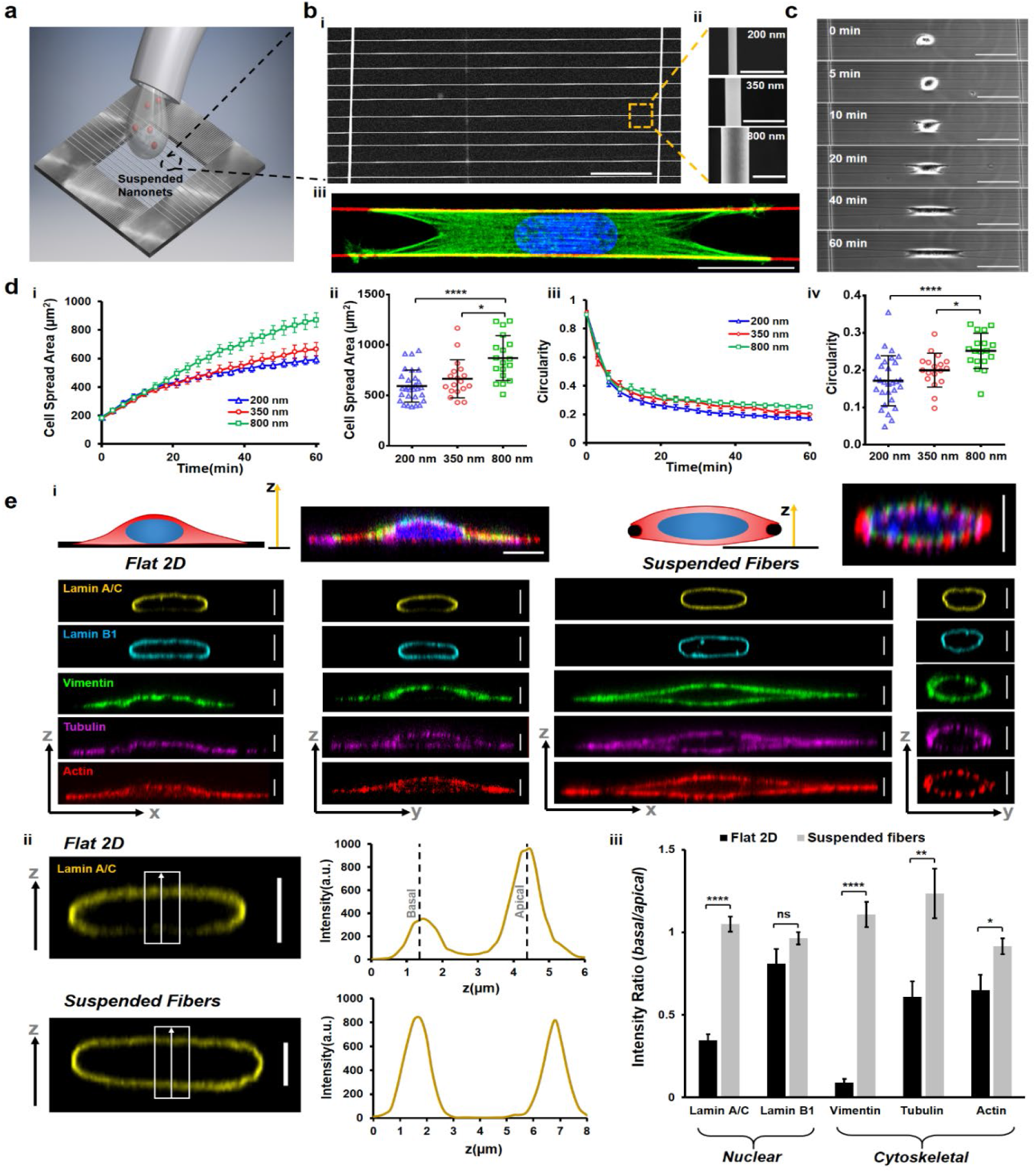
Cell spreading on suspended fiber networks: a) Schematic of assay used for studying cell spreading. Cell suspension is dropped on top of suspended nanonets coated with fibronectin and imaged in real-time. b) i) SEM image of suspended nanonet (scale bar, 50 μm) ii) SEM images of 3 different fiber diameters 200, 350 and 800 nm used for the study, (scale bar, 1 μm) and iii) Representative cell (stained for Actin-green, DAPI –blue, and fiber -red) on a 2-fiber (diameter: 350 nm) doublet (scale bar, 20 μm). c) Images from time lapse microscopy showing different phases of cell spreading on a 2-fiber doublet. (scale bar, 50 μm) d, i,iii) Temporal evolution of cell spreading area and circularity respectively (n=16-29 cells per category). d, ii, iv) Statistical comparison of cell spread area and circularity respectively after 1 hour of spreading e) i) Representative immunofluorescence images showing differences in localization of cytoskeletal elements and nuclear envelope proteins between Flat 2D and suspended nanonets. Cytoskeletal elements form a ‘caging’ structure surrounding the nucleus, while on 2D they ‘cap’ the nucleus from apical side (scale bars, 5 μm) and ii) Representative confocal side views (xz) along with intensity profiles along the z-direction, first and second intensity peaks correspond to basal and apical surfaces respectively, (scale bar, 5 μm) iii) Comparison of the basal and apical intensity for the different cytoskeletal and nuclear envelope proteins (n=10-11 per category) demonstrating intensity ratio values close to 1 for suspended nanonets (diameter: 200 nm) due to caging of nucleus, absent in cells on 2D. Images and data are shown for C2C12 myoblasts.

Next, we inquired if classic apical localization of cytoskeletal elements, described in studies on flat surfaces, extended to cells attached to suspended nanonets. On 2D substrates, nuclear shapes are shown to be regulated by cell boundary movements that involve both the actomyosin cytoskeleton’s contractility and the mechanical tension of the cell membrane ^[44,45]^. Khatau et al. identified the apical perinuclear actin cap as a primary regulator of the nuclear shape ^[46]^. In contrast, more recently, Ihalainen et al. demonstrated a differential localization^[47]^ of the nuclear lamins (particularly Lamin A/C) towards the apical nuclear envelope. Informed by these studies, we examined the localization of the major cytoskeletal elements (f-actin stress fibers, microtubules, and vimentin intermediate filaments) and nuclear lamins (A/C and B1) in cells attached to flat glass surfaces and on our suspended nanonets. We observed that the f-actin, microtubule, and intermediate filament cytoskeleton was highly aligned along the fiber axis in elongated cells compared to their counterparts on flat glass (**Figure S1 a,c**). Confocal microscopy side-views (xz and yz) at various stages during cell spreading revealed significant differences in the localization of the nuclear lamins and the overall organization of the cytoskeletal elements around the nucleus (**Figure 1e (i), Figure S2**). On flat 2D, our findings on apical-basal localization of Lamin A/C were in agreement with previously reported literature (*Intensity ratio*_*basal/apical*_ = *0.35±0.04*), but contrasted on suspended nanonets (*Intensity ratio*_*basal/apical*_ = *1.05±0.05*). In a similar manner, cells on flat glass displayed preferential enrichment of different cytoskeletal elements towards the apical side in a ‘capping’ manner (**Figure S2a**), with the strongest apical preference observed in the case of the vimentin intermediate filaments (*Intensity ratio*_*basal/apical*_ = *0.09±0.02*). Contrarily, the cytoskeletal network was nearly uniformly distributed around the nucleus in a ‘caging’ manner on suspended nanonets (intensity ratios ~1, **Figure 1e (iii)**). Overall, we conclude that cytoskeletal caging of the nucleus in suspended nanonets causes the uniform distribution of nuclear lamins, which is distinctly different than observations from 2D substrates.

### 2.2. Increase in cytoskeletal tension during cell spreading is fiber-diameter dependent

Since cells were achieving steady-state spreading fastest on the smallest diameters and suspended networks were causing cytoskeletal elements to cage the nucleus, we inquired about the role of focal adhesions and actin networks in establishing contractile forces. Visualizing cell-fiber adhesions through paxillin immunostaining at various time points (5, 10, 20, 40, and 60 min) during cell spreading revealed that focal adhesion (FA) clusters were formed along the entire cell length at early stages (t~10 min). In contrast, a preferential localization of the FA clusters occurred to the cell poles with increased spreading (**Figure 2a (i), Figure S3**). Normalized paxillin intensity taken along the cell length revealed two distinct peaks (**Figure 2a (ii)**) corresponding to the major FA clusters at either cell pole, consistent with our previous findings^[33,38,39]^. The transition of adhesion sites from being punctate along the entire cell body to localizing in major clusters at cell poles occurred by the 20-min time point (**Figure 2a (ii)**). A closer inspection of the adhesion distribution along the cell length at 60-min (**Figure 2a (iii)**) revealed smaller paxillin clusters distributed along the cell-fiber length, which became more prominent on larger diameter (800 nm) fibers (red arrowheads). FA cluster lengths were observed to grow in length with cell spread, with the largest 800 nm diameter fibers resulting in the longest cluster lengths at 60-min time point (**Figure 2a (iv, v)**).

**Figure 2:**
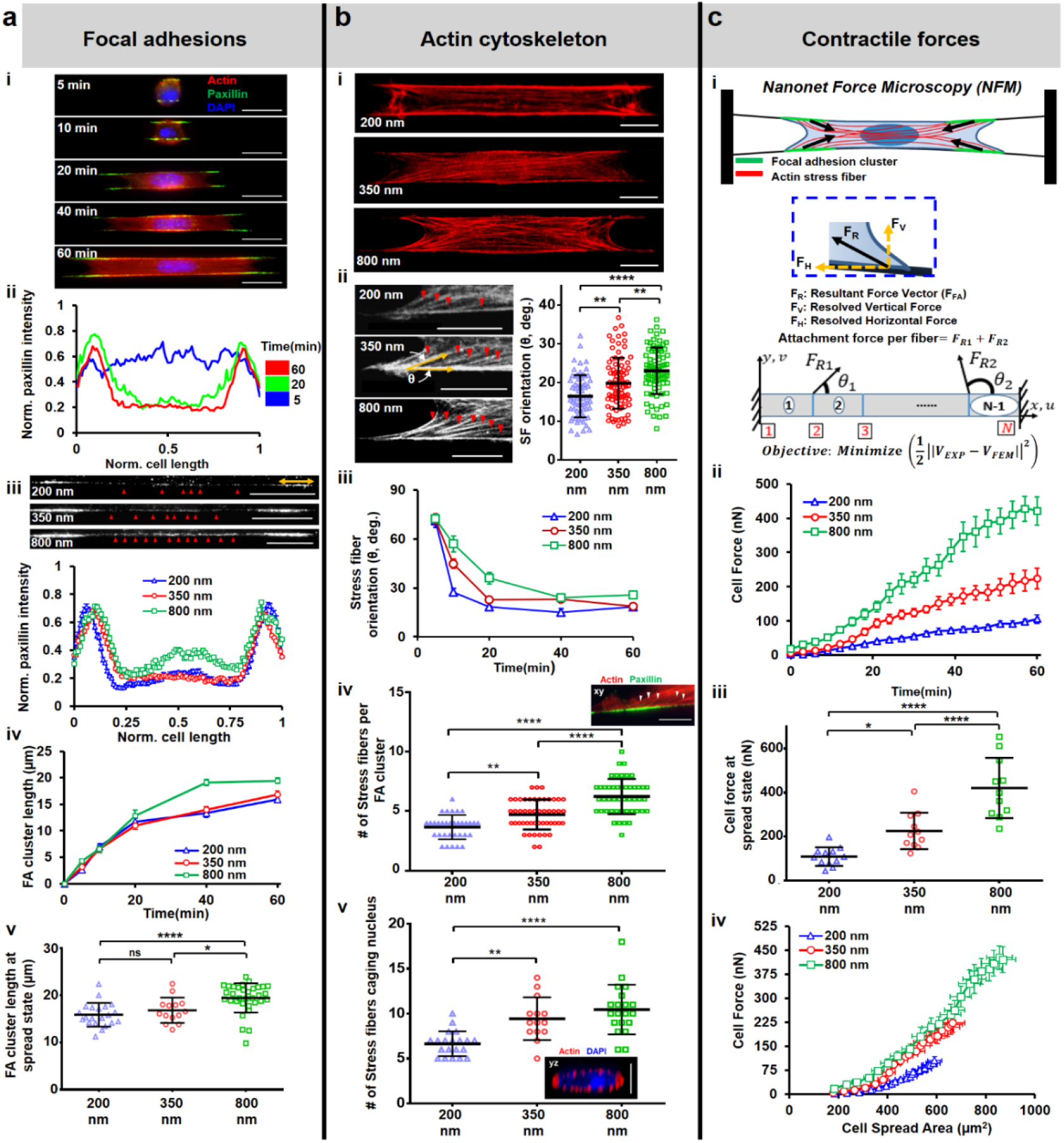
Cell contractility during spreading: a) Focal adhesion (FA) organization: i) Representative images showing cells stained for actin (red), paxillin (green) and DAPI (blue) at different timepoints during spreading (fiber diameter: 350 nm), (scale bars, 20 μm), ii) Normalized paxillin intensity (average of 9-12 profiles for each timepoint) showing spatiotemporal reorganization of focal adhesions as cells spread, iii) Differences in FA organization in clusters (yellow arrow) at the poles and along cell body-fiber length (red arrowheads) for different fiber diameters, (scale bars, 20 μm), iv) Temporal evolution of FA cluster lengths, and v) Comparison of FA cluster lengths at final spread state, n=21, 14 and 33, b) Actin cytoskeleton organization: i) Maximum intensity projections of actin cytoskeleton at spread state (60 min), (scale bars, 10 μm), ii) Stress fiber angle formed at the focal adhesion cluster zones, n=68, 88 and 80, (scale bars, 10 μm), iii) Transient evolution of orientation of the stress fibers during spreading, iv) number of major stress fibers originating from each FA clustering zone, n=33, 49 and 47 and v) number of major stress fibers caging the nucleus, n=20, 14 and 20. Inset are representative images of FA clustering zone, (scale bars, 10 μm) and nucleus caging with actin networks (scale bars, 5 μm), c) Contractile forces: i) Schematic of force measurement using Nanonet Force Microscopy (NFM) technique, ii) Transient cell attachment force evolution during spreading, (n=11-12 cells each diameter category), iii) Comparison of cell forces at spread state between the different fiber diameters, iv) Relationship between cell forces and cell spread area. Cell forces are computed from both fibers. Sample sizes are given for 200-200, 350-350, and 800-800 nm diameter doublets respectively. Images and data are shown for C2C12 myoblasts.

Our observations that FA clustering occurred in a fiber diameter-dependent manner suggested that fiber diameter might also influence the associated cytoskeletal tension. We immunostained for the contractile f-actin cytoskeleton (**Figure 2b (i)**) in cells at various time points (5, 10, 20, 40, and 60 min) and found that the average angle made by stress fibers anchored to the FA clustering sites at poles progressively decreased as cells spread, with the shallowest angle formed on 200 nm diameter fibers (**Figure 2b (ii, iii)**). We observed that the number of stress fibers originating at each FA cluster zone increased with fiber diameter, indicating increased contractility (**Figure 2b (iv)**). We also investigated the actin cytoskeletal organization in the perinuclear region since the actin stress fibers in this region directly affect nucleus shape regulation. Appearing as individual dots surrounding the nucleus, in confocal cross-sections (yz, **Figure 2b (v)**), we found that the number of stress fibers originating from individual FA clustering regions were less than those in the perinuclear region due to convergence of stress fibers emanating from FA clusters on either side, a behavior unique to anisotropically stretched cells in suspended nanonets (**Figure S4**).

Using NFM, we estimated the forces in cell spreading by monitoring the contractile inward deflection of fibers (**Figure 1c, 2c (i)**). Fiber deflections were subsequently converted into attachment forces at the individual FA clusters using inverse methods that minimize the error between computed and experimentally observed fiber deflections (Materials and Methods). The inputs to the computational framework include fiber properties and force vectors that originate from FA clustering zones (**Figure 2a (iii)**) and are directed along the average stress fiber orientation per fiber category (**Figure 2b (ii, iii)**). Consistent with our finding that the number of stress fibers increases with fiber diameter, the computed forces (F_cell/fiber_ = F_R1_ + F_R2_) also increase as cells spread and with an increase in diameter (**Figure 2c (ii, iii)**). We also plotted cell force against the cell spread area (**Figure 2c (iv)**). We found that for the same area across different diameters, cells attached to larger diameter fibers exerted significantly higher forces. Overall, our data suggest that increasing the fiber diameter causes cells to form larger focal adhesion clusters at the poles, leading to an increased number of actin stress fibers resulting in higher cell contractility.

### 2.3. Nuclear translocation of YAP is regulated by nuclear compression

Given our observations on the arrangement of F-actin networks caging the nucleus, we sought to investigate how the compression forces impacted the nucleus geometry and translocation of various transcription factors, including YAP (Yes-associated protein) ^[48]^, known for its central role in mechanotransduction. To determine nucleus geometry, we used confocal microscopy on DAPI-stained cells (**Figure 3a (i)**) that were fixed at various time points (5, 10, 20, 40, and 60 min). We employed three parameters: nucleus projected area (size in xy plane, top view), nucleus eccentricity (shape in xy plane), and the nucleus thickness (compression in xz plane, side view). Nucleus projected area increased steadily and reached an equilibrium value for all the fiber diameters tested (**Figure 3a (ii), Figure S5a**), with cells in 200 nm nanonets reaching stable areas the fastest. As cells spread, we observed the nuclei elongated (rise in eccentricity, **Figure S5b**). Not surprisingly, as the nuclei underwent compression from the near-spherical shape in rounded cells to the flattened ‘pancake’ shape (confocal side views, **Figure 3a (i)**), the nucleus thickness reduced significantly over time (**Figure 3a (ii)**), with the minimum thickness in cells attached to 800 nm nanonets (**Figure 3a (iii)**), suggesting that the cytoskeletal tension primarily drove the nuclear compression.

**Figure 3:**
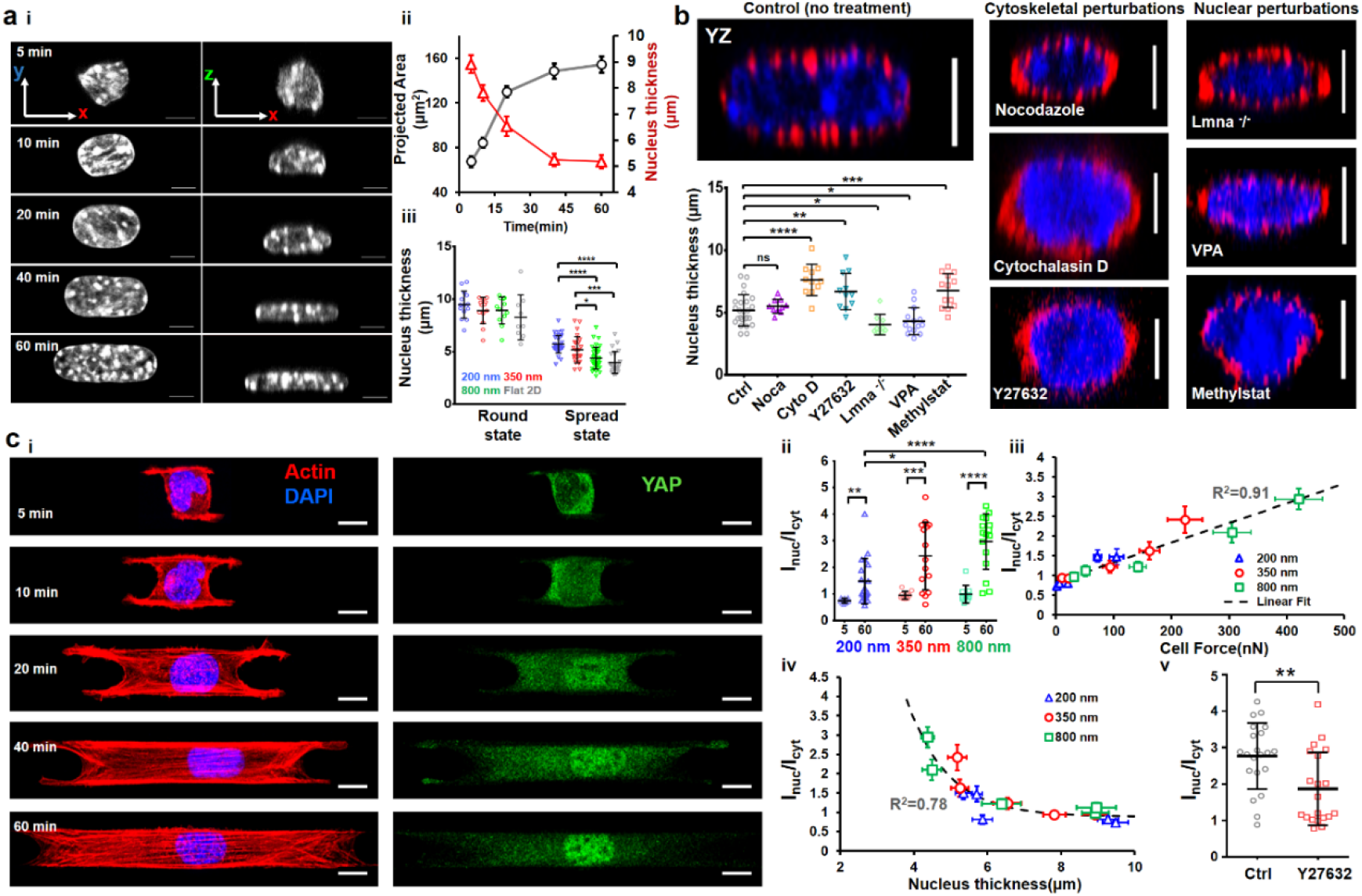
Nuclear compression and YAP localization during spreading: a) i) Representative images showing top view (xy) and side view (xz) of nuclei at different timepoints during cell spreading (fiber diameter: 350 nm), (scale bars, 5 μm), ii) Increase in nucleus projected area (xy) and decrease in nucleus thickness (z), indicating nucleus flattening with cell spreading, n=13-32 per timepoint iii) Nucleus thickness comparison between rounded (5 min) and spread (60 min) state, n=14, 13, 12 and 9 (for round state) and n=32, 25, 36 and 22 (for spread state) for 200 nm, 350 nm, 800 nm and Flat 2D respectively, b) Representative cross-section (yz) images of cells under various drug conditions and lamin A/C knockdown, n=25, 10, 11, 11, 9, 14 and 13 (scale bars, 5 μm), Comparison of nucleus thickness with control (no treatment), fiber diameter: 350 nm, c) Nuclear entry of YAP during cell spreading: i) Representative images at various timepoints during spreading, (scale bars, 10 μm) ii) Comparison of nuclear/cytoplasmic YAP between rounded (5 min) and spread state (60 min) for different fiber diameters, n=10-15 cells per timepoint and diameter category iii) Increase in nuclear YAP correlates with cell forces (iv) Increase in nuclear YAP correlates with nucleus thickness, and (v) Nuclear YAP localization decreases with loss of contractility through pharmacological inhibition, n=21 and 20 for control and Y27632 treatment. Fiber diameter: 800 nm. Images and data are shown for C2C12 myoblasts.

To test the relative contributions of the actin and microtubule cytoskeletons in regulating force-driven nuclear compression, we treated cells on nanonets with either cytochalasin D (2 μM) or nocodazole (1 μM). As expected, cytochalasin D treatment led to cells of reduced spread area (**Figure S6**) and disrupted actin cytoskeleton as evidenced by loss of stress fibers caging the nucleus (**Figure 3b**). Under these conditions, the nucleus thickness was significantly (~1.5x) higher than the control cells (no drugs). In contrast, disruption of the microtubule cytoskeleton did not significantly alter the nuclear compression levels. Next, we reduced cytoskeletal tension and actin stress fiber formation through the selective ROCK inhibitor Y27632 and observed nuclei of significantly larger thickness (**Figure 3b**), further elucidating the role of the actin cytoskeleton caging network in nucleus shape maintenance.

Informed by these findings, next, we investigated the role of the nuclear lamina in sustaining cytoskeletal forces. The nuclear lamina, composed of the nuclear lamins (A, C, B1, and B2), is a thin intermediate filament meshwork enveloping the nucleus, providing essential mechanical support. We generated C2C12 lamin A/C KD cells with wrinkled nucleus morphologies (**Figure S7**) consistent with observations from previous studies^[49]^. Fully spread Lamin KD cells had lower nucleus thickness than control, likely due to decreased nuclear stiffness^[15]^. Next, we altered the chromatin compaction levels by pre-treating cells with the histone deacetylase inhibitor (HDACi) valproic acid (VPA) used for chromatin decompaction and reducing nucleus stiffness or the histone demethylase inhibitor, Methylstat used for chromatin compaction and increasing nuclear stiffness^[8,50]^. Cells treated with VPA or Methylstat demonstrated significantly decreased and increased nucleus thickness, respectively, compared to control cells (**Figure 3b**).

Recent studies have shown how cytoskeletal-mediated compressive forces acting on the nucleus can stretch the nuclear membrane pores^[12]^. We wanted to inquire if the nuclear entry of YAP correlated with the fiber-diameter driven nuclear compression. Immunostaining for YAP (**Fig 3c (i)**) at various time points during cell spreading showed that YAP localization was primarily cytoplasmic (YAP intensity ratio between nucleus and cytoplasm I_nuc_/I_cyt_ <1) during early stages of cell spreading (5-10 min) but with an increase in cytoskeletal tension at later stages of cell spreading, a significant increase in the nuclear entry of YAP was observed (**Figure 3c (ii)**). Interestingly, we observed that the nuclear YAP translocation ratio was independent of cell shape but dependent upon force and nucleus thickness (**Figure 3c (iii, iv)**). Thus, reducing nuclear compression through contractility inhibition (Y27632 treatment) resulted in a significant decrease in the nuclear translocation of YAP (**Figure 3c (v)**).

### 2.4. Mismatch diameter fiber networks sculpt asymmetric nuclear shapes and invaginations

Cells in the native fibrous ECM can interact simultaneously with fibers of different diameter combinations. Hence, we interrogated force dynamics and YAP localization in mismatch diameter nanonets. To this end, we developed a fiber-spinning strategy using our non-electrospinning Spinneret-based Tunable Engineered Parameters (STEP^[51–53]^) platform to deposit nanonets in two mismatch-diameter combinations (2-fiber 200-800 nm doublets, and 3-fiber triplets (200-800-200 nm, and 800-200-800 nm). We reasoned that cell spreading on 200-800 nm doublets would proceed as described in Figure 1, with cells spreading faster on 200 nm side leading to trapezoidal-shaped cells with a longer base on 200 nm diameter side. Unexpectedly, we found symmetric spreading on both diameters (**Figure 4a, Movie S5**) but enhanced focal adhesion clustering (with respect to the 200 nm-200 nm counterparts) along the cell-fiber interface on the 200 nm fiber side (**Figure 4a (i, ii)**) that was similar to the focal adhesion clustering on 800 nm nanonets (**Figure 2a (iii)**). Increased cell adhesion sites on the 200 nm fiber side caused cells to spread more, have higher circularities, and exert larger forces on the 200 nm diameter fiber than their counterparts on 200-200 nm nanonets (**Figure 4a (iii-v)**).

**Figure 4:**
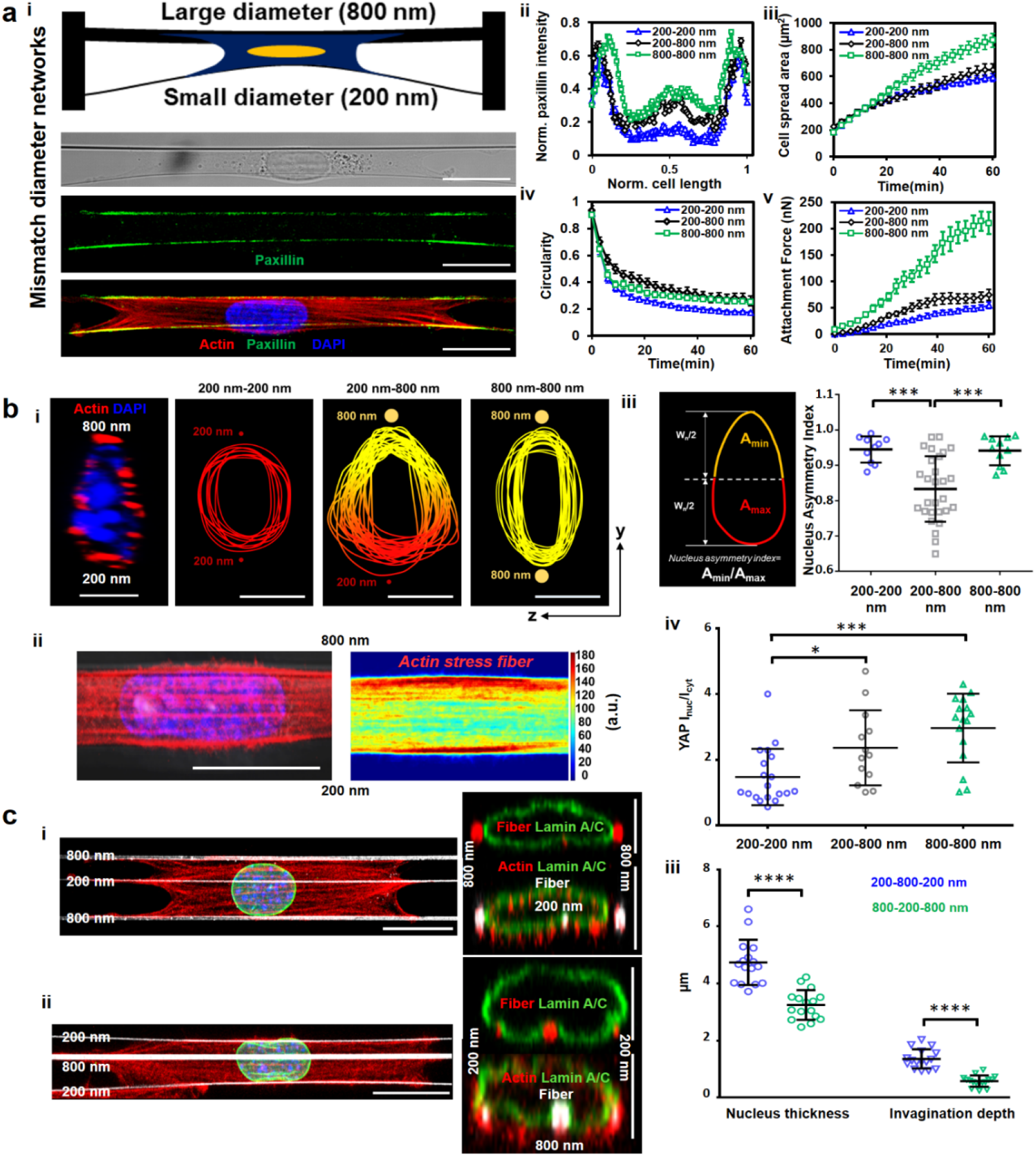
Sculpting nucleus shapes using mismatch diameter networks. a i) Cells attached to a small diameter (200 nm) fiber coupled on other side to a large diameter (800 nm) fiber, (scale bars, 20 μm) ii) Mismatch fiber combination leads to enhanced focal adhesion organization along smaller diameter fiber, n=10-14 per category, (iii) Increased cell spread area compared to 200-200 nm networks iv) Altered cell shape, n=12-17 per category, v) Force exertion on 200 nm diameter fiber in 200-800 nm mismatch configuration is higher than in 200-200 nm configuration, n=10-12 per category, b i) Mismatch (2-fiber, 200-800 nm) networks induce asymmetric (teardrop) nucleus shape, nuclear cross-sections (yz) perpendicular to cell length are used (scale bars, 5 μm), (ii) Teardrop shapes are formed due to higher density of actin stress fibers at the 800 nm diameter side of cell, n=17 cells, iii) Analysis of teardrop shapes using Nucleus Asymmetry Index, n=10,28 and 11 for 200-200, 200-800 and 800-800 combinations respectively, iv) nuclear/cytoplasmic YAP in mismatch diameter networks, n=19,13 and 16 for 200-200, 200-800 and 800-800 combinations respectively c) i,ii 3-fiber mismatch (800-200-800 nm and 200-800-200 nm) can induce ‘invaginations’ in nucleus (yz-cross sections, nuclear envelope stained for lamin A/C, green, fibers, red), second representative cross-section for each category, is included to demonstrate cytoskeletal caging, actin (red), Lamin A/C (green), fibers (white) and nucleus (blue), (scale bars, 10 μm for top views, 5 μm for cross-sectional views) iii) Comparison of the nucleus thickness and the depth of the invaginations show that 200-800-200 nm triplets have thicker nuclei and larger invaginations due to middle 800 nm diameter fiber compared to 800-200-800 category (n=16 for both categories). Images and data are shown for C2C12 myoblasts.

Intrigued by the altered force exerted by cells on mismatch nanonets, we next inquired if YAP localization inside the nucleus was also affected. First, we investigated the differences in nucleus cell shapes. We unexpectedly discovered asymmetric nuclear shapes (teardrop, confocal side view) with the tapered end towards the 800 nm diameter fiber (**Figure 4b (i)**). Analyzing the actin stress fiber distribution across the cells attached to mismatch diameter doublets revealed a higher density near the larger diameter fibers, potentially being the cause of increased nucleus compression (**Figure 4b (ii)**). To quantify the teardrop shapes, we defined the nucleus asymmetry index (NAI, **Figure 4b (iii),** Materials and Methods) as A_min_/A_max_, where A_max_ and A_min_ are the areas of the larger and smaller regions with respect to the nucleus mid-width line (**Figure 4b (iii)**) respectively. Cells adhering to mismatch doublets demonstrated significant asymmetry (NAI ~ 0.8, **Figure 4b (iii)**), compared with symmetric nuclei on 200-200 and 800-800 nm nanonets (NAI ~ 0.95). Furthermore, the nuclear localization of YAP was significantly enhanced in mismatch diameter fiber networks, as compared to the 200-200 nm, but less than 800-800 nm nanonets (**Figure 4b (iv)**).

Next, we extended the fiber-spinning strategy to generate precise 3-fiber nanonets (triplets) that provided the symmetric same-diameter outer fibers and a mismatch diameter inner fiber (200-800-200 nm and 800-200-800 nm, **Movie S6,** and **S7** respectively) that resulted in symmetric cell shapes (**Figure 4c (i,ii)**). However, we noted that the shape of the nucleus on outer fibers was flattened and at inner fiber had nucleus distortions (invaginations). Visualizing nuclei’s confocal cross-sections (yz), immunostained for Lamin A/C (nuclear envelope marker), revealed several interesting aspects of the overall nuclear geometry. First, the nucleus thickness observed on these mismatched 3-fiber nanonets demonstrated a similar trend compared to the same diameter 2-fiber nanonets. Nuclei in cells on 200-800-200 nm nanonets were significantly thicker than those on 800-200-800 nm, just as the nucleus thickness in 200-200 nm nanonets was higher than 800-800 nm nanonets (**Figure 4c (iii)**)). Second, in 3-fiber triplets, we observed the extent of invagination in the 200-800-200 nm combination was more significant due to lower force exertion of the outside 200 nm diameter fibers (**Figure 4c (iii)**). Overall, our data suggest that nucleus shape, YAP localization, and nucleus invaginations are regulated by the diameter combinations of fibers external to the cell.

### 2.5. Single fibers sculpt curvature-dependent nuclear invaginations

Our observation that single fibers induced nucleus invaginations running along the nuclei length prompted us to investigate the role of curvature in the physical indentation of the nucleus. To interrogate this behavior, we utilized a simplified model system of a single cell spreading on a single fiber (**Figure 5a**, **Movie S8**). Immunostaining for the focal adhesions (paxillin) revealed that similar to the parallel-cuboidal shape cells spread on 2-fiber nanonets, major focal adhesion clusters localized to the poles of the spindle shape cells (**Figure S8a (i)**). We also observed that majority of the actin stress fibers caged the nucleus (**Figure S8a (ii)**), causing local invaginations in the nucleus (**Figure 5b (i)**). We confirmed that the cell spreading and FA cluster formation was similar to our prior observations on 2-fiber nanonets (**Figure S8b (i, ii)**). Next, we varied the fiber diameter over a broad range (~150 nm (high curvature) to > 6,000 nm (low curvature) to understand the extent of fiber-induced local invaginations (**Figure 5b (i)**) in nuclei. We compared the radius of curvature (Ri, **Figure 5b (ii)**) of the nuclear membrane at the invagination site with the radius of the best-fit projected circle on the apical side of the nucleus (Ro). With increasing diameters, we found the curvature ratio Ri/Ro linearly scale with the fiber diameter (**Figure 5b (ii)**). We also found the nucleus aspect ratio reduced significantly with increased fiber diameter, indicating a change in nucleus shape from elongated on small diameter fibers to flattened on larger diameter fibers (**Figure 5b (iii)**). With the increase in diameters, we observed a drop in the cell aspect ratios (**Figure S8c**), with cells adhering to the 150 nm diameter fibers forming long protrusions and having the highest aspect ratios. Fluorescent labeling of the fiber (conjugated fibronectin, red) and the nuclear envelope (Lamin A/C, green) revealed the nucleus to be locally deformed at the location of fiber, thus allowing us to sculpt invaginations of varying sizes and shapes (**Figure 5c (i)**). We quantitated the effective shapes (**S**) of nuclear invaginations with a bell curve defined by 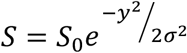, where S0 denotes the invagination depth (z-direction), and σ is related to the lateral spread (y-direction) of the invagination. We found that nuclear invaginations on lower diameter fibers of high curvatures were typically ~ 2 μm deep (S0) and narrow (σ ~ 0.4 μm). With an increase in fiber diameters, the size (depth and lateral spread) of these invaginations was found to increase (**Figure 5c (ii)**). However, the sharpness ratio (ratio of S0 and σ) was the highest for low diameter fibers, which decreased with an increase in diameter, indicating smoother nuclear deformations. Our data showed a strong linear fit between nucleus invagination shapes and fiber diameter for all tested diameters.

**Figure 5:**
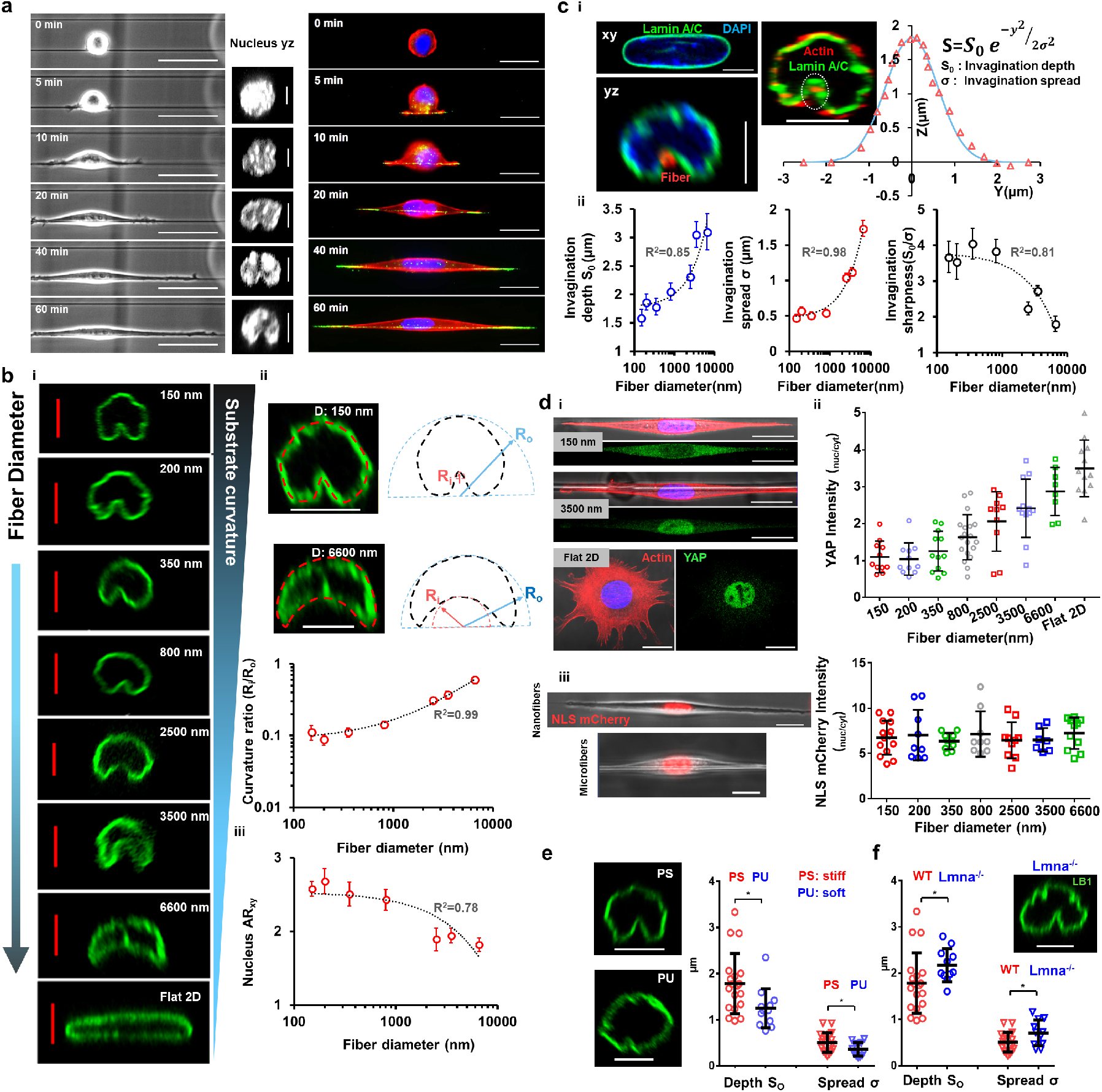
Sculpting nucleus invaginations using curvature of single fibers. a) i) Timelapse images showing different phases of cell spreading on a 1-fiber system causing cells to spread in elongated spindle shapes, (scale bars, 50 μm), Confocal side views of representative cells at various spreading time points demonstrate the fiber–mediated nuclear deformations (scale bars, 5 μm), cytoskeletal and focal adhesion arrangement in spindle-shaped cells at various timepoints, actin (red), paxillin (green) and nucleus (blue), (scale bars, 20 μm) b) i) Representative images showing effect of substrate curvature (fiber diameter) on the invagination shape/size, (scale bars, 5 μm) ii) representative images of the deformed nucleus shape on two diameters (150 and 6000 nm, scale bars: 5 μm) and methodology for comparing invagination shapes, Fiber diameter dependence of Curvature ratio: Radius of curvature at the invagination side (R_i_) divided by radius of curvature at the nucleus apical side (R_0_), n=12,10,17,10,10,9 and 9 respectively, (iii) Nucleus aspect ratio (top view, xy) as a function of fiber diameter, n=14,10,10,11,10,6 and 6 respectively c) i) Nucleus in spindle cells exhibit invaginations near sites of external fibers (red circle), Representative image showing cytoskeletal caging of nucleus in spindle cell, actin (red) and Lamin A/C (green), (scale bars, 5 μm), Invagination shape (shown in white circle) can be approximated with a bell curve, ii) invagination depth and spread increase with fiber diameter, while invagination sharpness decreases with increase in fiber diameter, n=9-19 cells per diameter category d) i) Representative stained images showing YAP localization in cells on nanofibers, microfibers and Flat 2D, (scale bars, 20 μm) ii) Comparison of YAP localization (I_nuc_/I_cyt_) in cells on different fiber diameters and Flat 2D, n=11,11,12,19,10,12,8 and 12 cells respectively, iii) Representative cells expressing NLS mCherry, showing primary nuclear localization of NLS on both nanofibers and microfibers, (scale bars, 20 μm), NLS localization is predominantly in the nucleus in cells attached to fibers of all diameters, n=7-14 cells for each diameter category e) influence of fiber stiffness in regulation of invagination size, n=19,12 for PS and PU respectively, (scale bars, 5 μm) and f) influence of impaired nuclear lamina on invagination size in cells attached to 350 nm diameter fibers, n=19,11 for control, Lamin A/C KD respectively (scale bars, 5 μm). All R2 values shown are calculated for linear fits. Sample sizes (4b (ii), 4c (ii), 4d (iv)) are given for 150 nm, 200 nm, 350 nm, 800 nm, 2500 nm, 3500 nm and 6600 nm respectively. Images and data are shown for C2C12 myoblasts.

Next, we wanted to investigate the effect of fiber diameter on the spatial localization of YAP. On the lower diameter nanofibers, cells primarily demonstrated a cytoplasmic YAP localization (**Figure 5d (i)**) with the nucleus/cytoplasmic intensity ratio less than ~ 1 (**Figure 5d (ii)**). However, with increased fiber diameters to micron-scale, we observed significantly enhanced YAP localization within the nucleus (**Figure 5d (ii)**). Interestingly, despite various levels of nuclear deformations in both nanofibers and microfibers, the nuclei demonstrated no signs of rupture, as confirmed by no appreciable leakage of NLS (nuclear localization sequence) in cells expressing NLS mCherry (**Figure 5d (iii), Figure S9, Movie S9**). Quantification of NLS localization revealed an average nuclear/cytoplasmic intensity ratio > 6 for all fiber diameters, demonstrating NLS to be primarily localized to the nucleus.

Next, to investigate the effect of material stiffness, independent of fiber curvature, we used polyurethane (E: 10-100 MPa) and polystyrene (E: 1-3 GPa) fibers of ~350 nm diameters. We observed a significant decrease (**Figure 5e**) in the size of the nuclear invaginations in softer polyurethane fibers. We also inquired about the role of lamin A/C in nucleus invaginations and found that impairing the nuclear envelope caused by the knockdown of lamin A/C resulted in significantly larger nuclear deformations (**Figure 5f**). Inhibiting actin-based contractility through Y27632 treatment resulted in thicker nuclei with significantly smaller nuclear invaginations (**Figure S8b (iv)**).

### 2.6. Multiple fibers of the same diameter sculpt nucleus shapes

Since single fibers were causing significant invaginations in the nuclei, we inquired if multi-fiber networks (≥3) of the same diameter caused multiple similar-sized invaginations. We deposited 350 nm diameter fibers at low (~3 μm) inter-fiber spacing (**Figure 6a (i)** images shown for single-cell attached to five and eight fibers). We found that cell and nucleus aspect ratios (at the spread state) were reduced with the increase in the number of fibers (**Figure 6a (ii), Movie S10** for three fibers, and **S11** for 7 seven fibers). Also, the increased compression mediated by the cytoskeletal caging caused a significant reduction in nucleus thickness and invagination depth (**Figure 6a (iii)**). Additionally, these deformations failed to generate ruptures in the nucleus, as confirmed by the spatial localization analysis of NLS (**Figure S9**).

**Figure 6.**
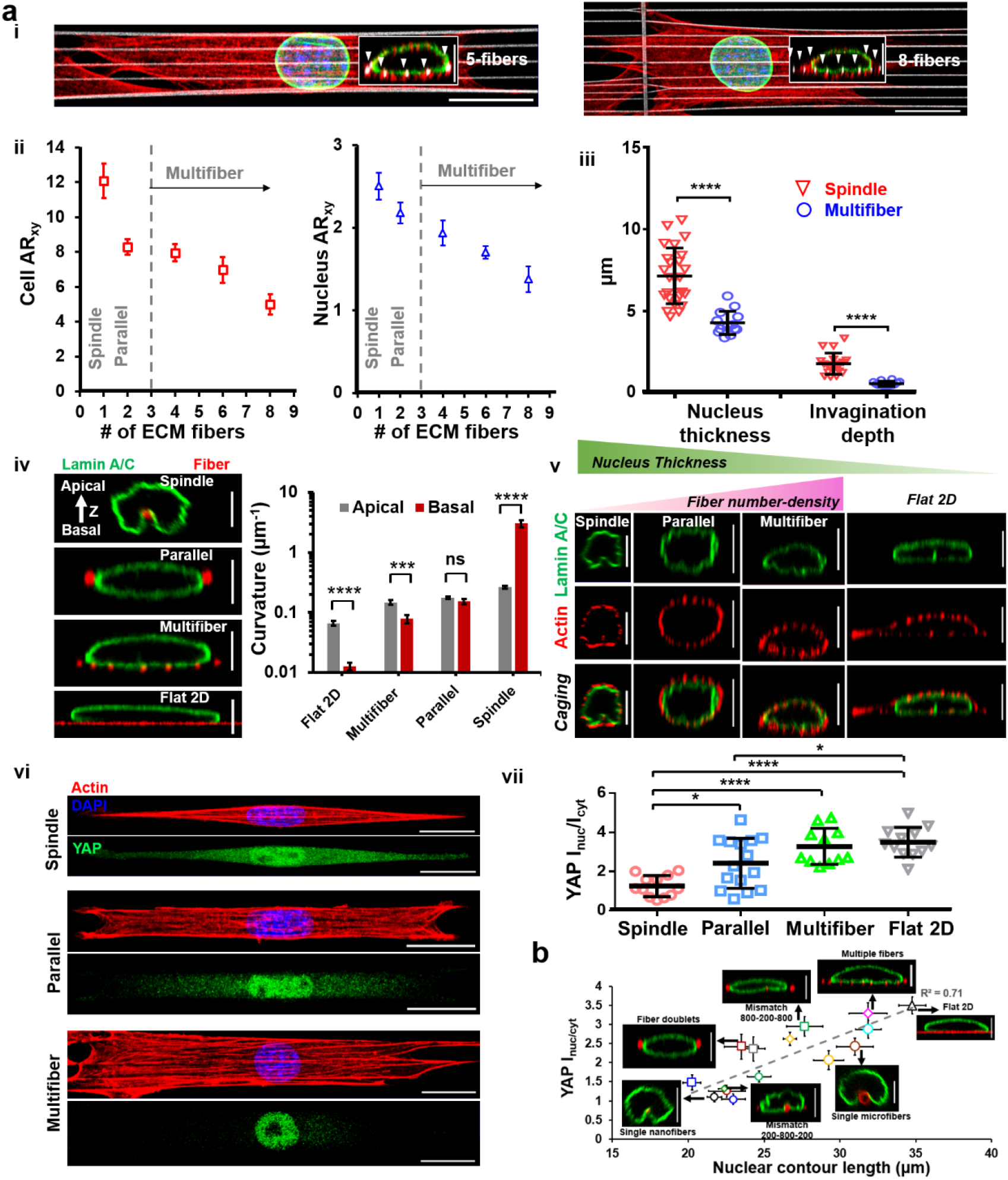
Nuclear geometry and YAP localization in cells attached to multiple fibers. a) All figures are for 350 nm diameter fibers. i) Representative images of cells attached to 5 and 8 fibers respectively, stained for actin (red), Lamin A/C (green) and nucleus (blue). Fibers are coated with rhodamine fibronectin (pseudo-colored white), Insets are cross sections (yz), showing the cytoskeletal caging of the nucleus with fibers identified with white arrowheads (scale bars, 20 μm and 5 μm for main and inset images respectively), ii) comparison of the cell and nuclear aspect ratio (xy top view) as a function of the number of interacting fibers, n=10, 24, 6, 7 and 5 iii) nucleus thickness (n=25,14) and invagination depth (n=19,14) in spindle (1 fiber) versus multifiber (≥ 3 fibers) systems, and iv) Representative nuclear cross sections (yz) demonstrating shape sculpting over different substrates, Analysis of curvatures at the apical and basal side of the nucleus for the different substrates, note, only curvature magnitude is considered here, spindle basal curvature being inward, is of opposite sign as compared to all other categories, n=11, 16, 11 and 18 for Flat 2D, multifiber, parallel (2-fiber) and spindle (1-fiber) respectively, v) Representative images showing the caging effect and nuclear compression as a function of fiber-number density, Flat 2D is shown for reference, actin (red), Lamin A/C (green) (scale bars, 5 μm), vi, vii) Comparison of YAP localization between spindle, 2-fiber, multifiber systems and flat 2D, n=12, 15, 11 and 12 respectively, Representative cells are stained for actin (red), YAP (green) and nucleus (blue), (scale bars, 20 μm). b) YAP localization across single, 2 fiber and multifiber configurations of all diameters and interfiber spacing in the study showing positive correlation between YAP localization and increase in contour length (n=8-16 cells for each substrate category), small and large circles represent nanofibers and microfibers, squares represent fiber doublets (same diameter and mismatch), polygons represent 3-fiber mismatch and multifiber configurations, Flat 2D is marked with triangle. Images and data are shown for C2C12 myoblasta.

We also quantified the shapes of nuclei in cells attached to flat 2D, 1-fiber, two- fiber, and multiple fiber systems by quantifying the curvature of the nuclear envelope on either side (**Figure 6a (iv)**). Consistent with previous literature^[47,54]^, our results indicate that the nucleus is mostly pancake-shaped on flat continuous surfaces, which differs from cells on suspended 2-fiber parallel doublets, where the nucleus is mostly ellipsoid shaped, with equivalent curvatures on both sides. In multifiber networks, we found the nucleus shape to have a flatter surface (low curvature) at the basal side, primarily due to cell attachment to multiple underlying fibers. In comparison, the fiber-induced invaginations in the spindle-shaped cells result in a much sharper curvature (inward) at the basal side. Concurrent with our previous findings that increasing the number of fibers results in increased cell contractility^[42]^, we found that for the same diameter, increasing the number of fibers results in a decrease in nucleus thickness (**Figure 6a (v)**). Interestingly, enhanced nuclear compression caused by cell attachment to multiple fibers led to a significant increase in YAP’s nuclear translocation compared to spindle and parallel-cuboidal cells attached to single or two fibers of the same diameter, respectively (Figure 6a (vi-vii)). To understand how different fiber configurations led to the altered nuclear entry of YAP, we computed the stretching of the nuclear contour (yz cross-section) due to local invaginations (single fibers) or compression (fiber doublets), or both (multiple fibers). We observed a distinct increase in the contour length in cells attached to micron-scale fibers or multiple fibers. In general, nuclear YAP localization for different fiber configurations and Flat 2D demonstrated a strong dependence on the nuclear contour length (R^2^=0.71, **Figure 6b**).

Overall, our observations show that the organization of contractile f-actin network in spindle cells on single fibers induces nucleus invaginations in a diameter dependent manner, while the same f-actin network in cells attached to multiple fibers leads to higher levels of nuclear compression but reduced local invaginations and causes distinct YAP translocation to the nucleus. Irrespective of the shape, sharpness, and number of invaginations, we find that nuclei of cells attached to fibrous matrices remain rupture-free.

## 3. Discussion

Nucleus shapes have been studied in a wide range of *in vitro* settings, but the role of physiological fibrous ECM such as those found in skeletal muscle tissue and around metastatic tumors ^[26,27,55]^, in controlling nucleus shapes remain poorly described. Fibrous ECM environments are composed of a mix of diameters (individual nanoscale fibrils that combine into larger hundreds of nanometer or micron-scale bundles) distributed in a wide range of orientations and inter-fiber spacing. In ECM regions with a large number of contact sites (small pore size), cells sense confinement, and with few contact points (large pore size), cells make contact with only single fibers ^[56–58]^. Cell contractility is altered with differing contact points across multiple diameters ^[38]^, resulting in changes in the overall cell ^[37,39]^ and nucleus shape. Here, we explore the mechanosensitivity of cells to fibrous microenvironments using nanofiber networks of precisely controlled geometries and fiber diameters. Our method of precisely controlling the external fibrous environment led to exquisite sculpting of the nucleus with and without invaginations across multiple cell lines (C2C12 mouse myoblasts, HT1080 human fibrosarcoma, and HeLa cells, Figure S10). Irrespective of the shape or sharpness of the invaginations, we found nuclei to remain rupture-free and fiber-diameter driven contractility and network organization to regulate translocation of mechanosensitive YAP to the nucleus.

To understand fiber diameter-driven contractility and associated nucleus shape changes, we estimated the increase in contractility as rounded cells spread along the fiber doublets. While the cell spreading behavior on fibers was similar to reported flat 2D behavior ^[59]^, the spreading rates were fiber diameter dependent, with cells attached to thinnest fibers achieving steady-state spread areas the earliest. In the spread state, although actin, microtubules, and vimentin localize at the apical side in cells on 2D, we found them to *cage* the nucleus with no apical-basal preference in suspended cells. Even in cells spreading on single or multiple fibers, where cell contact regions with fibers act as pseudo-basal regions, we observed a uniform distribution of Lamin A/C expression (**Figure S8d**). In all diameter combinations, focal adhesions were preferentially localized to the poles as cells spread on the fibers, consistent with their localization pattern during spreading at the peripheries in cells on flat 2D and the poles in cells on micropatterned substrates ^[60]^. The clustering of focal adhesions to the poles occurred early in the cell spreading phase (~20 min), resembling those observed in 3D cell-derived matrices (37, 38). Additionally, our observations that the lengths of FA clusters at the poles and the number of focal adhesions along the cell-fiber contact length increased with fiber diameter may potentially explain the slower spreading observed on larger diameter fibers.

Cell contractility increased with diameter, and we compared forces, estimated using Nanonet Force Microscopy (NFM) with other force measurement techniques using two metrics: traction stress and tension per stress fiber. Assuming that the forces are uniformly distributed over the entire length of FA cluster, then the average stress per FA cluster (Force/ FA cluster_area_, where FA cluster_area_ = FA cluster_length_ * Diameter_fiber_) was found to be ~8, 9, and 7 nN/μm^2^ for 200-200, 350-350, and 800-800 nm doublets, respectively, which is similar to the reported values of stress per focal adhesion in the literature (~6 nN/μm^2^) ^[63,64]^. We approximated the tension in individual stress fibers by assuming their arrangement to be similar to be mechanical springs in parallel. Dividing the force per FA cluster at spread state (60 min time point) with the number of stress fibers associated with each cluster, we obtained the tension in each stress fiber to be ~7, 11, and 15 nN for 200-200, 350-350, and 800-800 nm doublets, respectively, which agree with reported values (2-10 nN) (41, 42). The ability to estimate spreading forces allowed us to examine the extent of nucleus compression as a function of fiber diameter. Since 800-800 nm doublets were exerting the highest forces, not surprisingly, the nuclei in these networks had the highest compression that matched the thickness of nuclei on flat 2D ^[54,67]^. Loss of actin tension via cell treatment with cytochalasin D alleviated nucleus compression, a behavior consistent with previous findings ^[68]^. In contrast, the loss of microtubules via nocodazole treatment did not affect the nucleus compression. On the other hand, reduction in stiffness of the nucleus and lamin A/C KD resulted in increased compression while stiffening the nucleus reduced nucleus compression. Previous studies have shown that nucleus flattening causes stretching of the nuclear pore complexes ^[12]^ (NPCs), which lead to the nucleo-cytoplasmic shuttling of various transcription factors, including YAP/TAZ ^[12]^ and HDAC3 ^[14]^. Across the tested diameters, our studies concluded that YAP localization increased with cell spreading due to increasing contractility. Importantly, our studies showed that for similar levels of contractility but different spread areas, YAP translocation remained similar.

We also generated *in vitro* fibrous environments of a mix of diameter combinations to mimic ECM heterogeneous fibrous environments. First, we generated 200-800 nm fiber diameter mismatch doublets to study altered cell spreading, contractility, and nucleus shape. Unlike FA clustering patterns observed during cell spreading on homogenous doublets (200-200 or 800-800 nm), unexpectedly, we found a higher density of focal adhesion sites on the 200 nm diameter that matched the adhesion sites’ pattern on the 800 nm diameter fiber, resulting in delayed cell spreading. Moreover, the 200 nm diameter fibers deflected more than the corresponding deflection in a 200-200 nm homogenous doublet, resulting in increased contractility. Confocal side views (yz) in mismatch doublets showed the nuclei to have a remarkable teardrop shape (~60% cases), with the broad end towards the 200 nm diameter. Intensity analysis of f-actin stress fiber distribution across the cell body in mismatch doublets revealed a higher density of stress fibers towards the 800 nm diameter fiber, thus contributing to the increased compression of the nucleus. We then generated triplets composed of two same diameter outer fibers and a third middle fiber of different diameters (200-800-200 nm, and 800-200-800 nm). We observed the 800 nm diameter fiber to induce large invaginations in the nucleus in 200-800-200 nm triplets, consistent with the finding that the outer larger diameter pairs caused significant nucleus compression due to higher contractility. The effect of middle fiber in either of these triplets did not affect cell spreading behavior.

Nucleus invaginations have been previously reported using micropillars at sites of contact ^[69,70]^. In contrast, here, we demonstrate that the invagination in the nucleus occurs across the entire length of the nucleus. Naturally, we inquired if these invaginations’ size depended on the fiber diameter. Our earlier work has shown that cells attached to < ~100 nm diameter fibers remain rounded, while those attached to larger diameters form elongated spindle shapes^[34]^. Thus, we chose the fiber diameters to range from 150-6600 nm. Immunostaining for actin and paxillin revealed that actin stress fibers connecting the FA clustering zones straddled the nucleus, thus causing compression of the nucleus and invaginations. The high curvature (low diameter) fibers caused the sharpest invaginations (*S*_*0*_/*σ*), which transitioned to smooth, wider invaginations with an increase in diameter. Despite such extreme nuclear invaginations during cell spreading, no nuclear rupture events were detected (Movie S9). Recently, single migrating cells have been shown to undergo nucleus rupture when they are subjected to confinement of ~3 μm ^[71]^. On the other hand, our studies were able to get the nucleus to compress to a height of ~4 μm. Under these conditions, we show that while fiber curvature and compression directly contribute to YAP localization, they are insufficient to cause nucleus rupture. Increasing the number of fibers of the same diameter (350 nm) further decreased the nucleus’ height due to increased contractility, resulting in reduced invagination depth compared to a single 350 nm diameter fiber. Contour length, a measure of the stretching of the nucleus, shows a positive correlation with YAP localization across the different diameters and interfiber spacing tested in our study (**Figure 6b**). In all the combinations tested by us, nuclei remained rupture-free during cell spreading.

## 4. Conclusion

In conclusion, we demonstrate sculpting of the nucleus and their functional response using in vivo mimicking fibrous environments through regulation of cell contractility in a fiber-diameter and number-density manner. The nuclei remain robust mechanical organelles capable of withstanding extensive shape changes and invaginations without undergoing rupture. We envision that control of nuclear 3D shape in these environments will yield new fundamental insights on the ECM-mediated alterations in the sub-nuclear chromatin organization and overall gene expression during cancer metastasis, wound healing, cell differentiation, and myogenesis.

## 5. Experimental Section/Methods

### 5.1 Fabrication of nanofiber networks

Using the previously reported non-electrospinning STEP technique^[51–53]^, suspended fiber nanonets (horizontal arrays of densely spaced (~12 μm) nanofibers of differing diameters (200nm, 350nm, 800nm) deposited on widely spaced (~ 250 μm) vertical support fibers (~2 μm diameter), were manufactured from solutions of polystyrene (MW: 2,000,000 g/mol; Category No. 829; Scientific Polymer Products, Ontario, NY, USA) was dissolved in xylene (X5-500; Thermo Fisher Scientific, Waltham, MA, USA) in 7-13 wt% solutions. Micron scale (≥ 2 μm) fibers were manufactured from 2-5 wt% of high molecular weight polystyrene (MW: 15,000,000 g/mol, Agilent Technologies, Santa Clara, CA, USA). Briefly, the polymeric solutions was extruded through a micropipette (inside diameter: 100 μm; Jensen Global, Santa Barbara, CA, USA) for deposition of aligned fibers on a hollow substrate. Measured (using scanning electron microscopy images) diameters for the 150 nm, 200 nm, 350 nm, 800 nm, 2.5 μm, 3.5 μm and 6.6 μm diameter categories are 153 ± 1.2 nm, 206 ± 2.6 nm, 361 ± 4.5 nm, 808 ± 7.6 nm, 2.50 ± 0.02 μm, 3.48 ± 0.09 μm and 6.62 ± 0.17 μm (values are shown as Mean ± SEM, n=37, 59, 90, 92, 28, 21 and 25 respectively). For the 350nm diameter category, fiber networks with varying spacing (3-25) μm were used. All fiber networks were crosslinked at intersection points using a custom fusing chamber, to create fixed-fixed boundary conditions.

### 5.2 Cell culturing and experimental procedure

C2C12 mouse myoblasts (ATCC), HT1080 human fibrosarcoma and HeLa cells were cultured in Dulbecco’s modified Eagle’s medium (Invitrogen, Carlsbad, CA, USA) supplemented with 10% fetal bovine serum (Gibco, Thermo Fisher Scientific) in T25 flasks (Corning, Corning, NY, USA) kept at 37°C and 5% CO_2_ in a humidified incubator. For imaging cell spreading dynamics scaffolds containing fiber nanonets were mounted in glass-bottom single well plate. For immunofluorescent staining experiments, cell seeding was performed on fiber scaffolds mounted on glass-bottom six-well plates (Cellvis, Mountain View, CA, USA). Prior to experimentation, fibers were sterilized in 70% ethanol for 10 minutes and functionalized for 1 hour under incubation at 37°C using 4 μg/mL fibronectin in PBS (Invitrogen, Carlsbad, CA, USA). For fluorescent labelling of fibers, rhodamine conjugated fibronectin (Cytoskeleton Inc., Denver, CO, USA) was used with the same concentration and incubation time. Before cell seeding, the test platform was moved into an AxioObserver microscope under incubation conditions of 37°C and 5% CO_2_ (Zeiss, Oberkochen, Germany). Without disturbing the closed environment, a droplet (~100 μL) of cell suspension was deposited on the fiber nanonets to begin the experiment.

### 5.3 Live imaging

Time-lapse optical imaging was started immediately before introducing the cell suspension to the fibers. Imaging was performed at 20x 0.8 NA objective in a Zeiss AxioObxerver Z1 microscope with an interval of 1-3 minutes for 1 hour. C2C12 or HT1080 cells expressing NLS (nuclear localization signal) mCherry were imaged every 3 min with a TRITC filter set.

### 5.4 Immunofluorescent staining and imaging

Cells were fixed with 4% paraformaldehyde for 15 minutes at various timepoints (5, 10, 20, 40 and 60 minutes), following initial cell fiber contact. Cells were then permeabilized with a 0.1% Triton X-100 solution, washed in PBS twice and blocked with 5% goat serum (Invitrogen, Grand Island, NY) for 30 minutes. Primary antibodies, diluted in an antibody dilution buffer consisting of PBS with 1% Bovine Serum Albumin and Triton-X 100, were added to the fixed cells and kept either i) overnight at 4° C / ii) at room temperature for 3 hours / iii) at 37°C for 2 hours. Diluted secondary antibodies, along with the conjugated Phalloidin-TRITC (Santa Cruz Biotechnology, Dallas, TX, USA) or Alexa Fluor 647 Phalloidin (Invitrogen) diluted in 1:80 ratio, were subsequently added and stored in a dark place for 45 minutes. Following a 3 times PBS wash, DAPI (4′,6-diamidimo-2-phenylindole) or Hoechst 33342 (Thermo Fisher Scientific) was added for 5 minutes to stain the cell nuclei. Primary antibodies include Anti-Vimentin antibody (1:250, rabbit monoclonal, EPR3776, Abcam), Anti-phospho-Paxillin (1:100, rabbit polyclonal, pTyr31, Invitrogen), Anti-beta tubulin (1:500, mouse monoclonal, 2 28 33, Invitrogen), Anti-Lamin A/C (1:1000, mouse monoclonal, sc-376248, Santa Cruz Biotechnology), Anti-Lamin B1 (1:500, mouse monoclonal, sc-374015, Santa Cruz Biotechnology) and Anti-YAP (1:100, mouse monoclonal, sc-101199, Santa Cruz Biotechnology). Secondary antibodies include Goat anti-Rabbit IgG Alexa Fluor 488 (1:200, Invitrogen), Goat anti-mouse IgG Alexa Fluor 488 (1:200-1:1000, Invitrogen) and Goat anti-mouse IgG Alexa Fluor 647 secondary antibody (1:500, Invitrogen). Images were taken using an inverted Zeiss microscope using a 63x objective (NA 1.20, water immersion). Confocal microscopy was performed using a laser scanning confocal microscope (LSM 880, Carl Zeiss Inc.) and images were obtained using a 63x 1.15 NA water immersion objective. Z-stacks were taken with slice thicknesses ranging between 0.3-0.5 μm. Z-stack images were processed in the Zen Blue software (Carl Zeiss Inc).

### 5.5 Generation of knockdown cell lines

Lmna-KD were created using shRNA encoded on pLKO.1 puro plasmid (Addgene #8453) and introduced into C2C12 with 2^nd^ generation lentivirus. Sequences for shRNA was obtained from Broad Institute Genetic Perturbation Platform. The sequence used for this study is GCGGCTTGTGGAGATCGATAA. Viral particles were produced in HEK293T following calcium phosphate transfection using 4.8 μg, 30 μg and 33.6 μg of VSVG (Addgene #14888), psPAX2 (Addgene #12260) and pLKO.1 puro Lmna-shRNA plasmids respectively. After 48 hrs, supernatant containing viral particles were concentrated 100x following 2-hr ultracentrifugation 50,000 g and 4°C. Cells were transduced in full growth media (DMEM supplemented with 10% FBS) containing viral particles for 48 hrs and verified (**Figure S11**) via Western blot using anti-Lmna/c mouse mAb (Cell Signaling Technology #4777).

### 5.6 Pharmacological inhibition

For cytoskeletal inhibition, cells in suspension were incubated with 2 μM cytochalasin D (actin disruption) (Fischer Scientific), 1 μM nocodazole (microtubule disruption) (Sigma Aldrich) and 10-20 μM Y27632 (ROCK inhibition) (Hellobio, Princeton, NJ, USA) for 30 minutes −1 h. For chromatin compaction /decompaction cells were pre-treated with histone demethylase inhibitor (Methylstat, 5 μM, 48 h incubation) (Sigma Aldrich, St. Louis, MO, USA) and histone deacetylase inhibitor valproic acid (VPA, 2 mM, 24 h incubation) (Sigma Aldrich) respectively. At least N=2 replicates were performed for each drug condition.

### 5.7 Analysis of shape metrics

Cells in suspension demonstrate a rounded morphology which evolves as they attach and spread on a substrate. Cell circularity is defined by the relation:

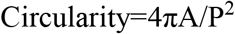

where A is spread area (μm^2^) and P is perimeter (μm). A perfect circle results in a circularity value of 1.0. As the shape elongates, this value approaches 0. Polarized cells were manually outlined in ImageJ and the aspect ratio was quantified using the Bounding Rectangle Function.

### 5.8 Analysis of focal adhesion clusters and actin stress fibers

Consistent with our previously reported convention, we quantified the length of the focal adhesion (FA) cluster as the longest continuous length of immunostained paxillin. Intensity profiles for paxillin were generated by performing line scans along the cell-fiber contact region. For averaging purposes, profiles were normalized in 2 steps, i) Intensity (plotted along y-axis) was normalized with respect to peak intensity of the corresponding profile, ii) Cell length (plotted along x-axis) was normalized with respect to the net length (measured from end-to-end) of the particular cell. Intensity profiles were subsequently averaged using a custom MATLAB routine, using a bin width of 0.01. Maximum intensity projections of phalloidin stained cells were utilized to count the number of major actin stress fibers emanating from each FA cluster and in the perinuclear region. Orientation of the actin stress fibers were measured with respect to the undeflected nanofiber orientation. For average actin localization in **Figure 4b**, images of perinuclear regions (17 cells) were averaged using a custom MATLAB routine, and the average image is represented as an intensity heatmap.

### 5.9 Analysis of nucleus shapes and deformations

Confocal z-stacks of DAPI/Hoechst/Lamin stained nuclei were processed in either ImageJ (NIH; https://imagej.nih.gov/ij/) or Zen Blue (Carl Zeiss Inc). Nucleus projected area was calculated from the top (xy) view at the equatorial plane (plane of the nanonets). The shape (xy) of the nucleus was approximated as an ellipse and quantified by the ellipse eccentricity. Eccentricity= (1-b^2^/a^2^)^0.5^ where 2a and 2b represent the shape of the major and minor axis of the ellipse respectively. The eccentricity value of a perfectly symmetrical circle is 0, while the value increases to 1, with increasing elongation of the ellipse. Nucleus aspect ratio was measured by manually outlining and using the Bounding Rectangle Function in ImageJ. Nucleus thickness was measured from the orthogonal side views (xz or yz) generated with the *Ortho* function in Zen Blue, and the maximum z-dimension was considered as the thickness of the nucleus. To quantify nuclear shapes in ‘mismatch’ diameter fiber doublets, cross-sectional side views (yz) were generated using *Ortho* function in Zen Blue. A line is then drawn through the nucleus mid-width as shown in (**Figure 4b (iii)**) and the area of the half cross-sections were measured and designated as A_max_ and A_min_ based on larger and smaller area respectively. Nucleus asymmetry index (NAI) is defined by A_min_/A_max_. Thus symmetric nuclei will have a NAI of ~1, while teardrop shaped nuclei will have NAI < 1.

To quantify nuclear invaginations in spindle shaped cells attached to single fibers, the deformed region of the nuclear envelope (yz cross-sectional view) was manually outlined in ImageJ and a bell curve (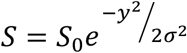, **Figure 5c**) was fitted to the manual trace using a custom MATLAB (https://www.mathworks.com/) code to evaluate the individual parameters S0 (invagination depth) and σ (invagination spread). To quantify the level of curvature of the nuclear surface, a best-fit circle was placed at the apical and basal surface of the cross-sectional view (yz) of the nucleus, in the Zen Blue software. The curvature was defined as the inverse of the radius of this best-fit circle.

### 5.10 Fluorescent Intensity analysis

To compare the basal and apical localization of cytoskeletal and nuclear envelope components on fiber doublets, the orthogonal side views of the respective channels were first generated in Zen Blue. Rectangular intensity profiles with a 5 μm width were computed at the cell/nucleus mid-width (**Figure 1e**). The peak intensities at the apical and basal side were subsequently extracted to compute the intensity ratio between basal and apical side. To quantify the YAP/NLS localization within cells, cells and nuclei were first manually outlined in Zen Blue to get the mean YAP/NLS intensity within the whole cell (I_cell_) and inside the nucleus (I_nuc_) and the corresponding projected areas (A_cell_ and A_nuc_). The average intensity within the cytoplasm (projected area: A_cyt_) is indirectly quantified from the following relations:

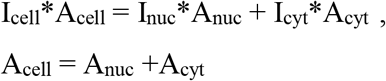

The intensity ratio I_nuc/cyt_ is quantified by the ratio of the average intensities within the nucleus (I_nuc_) and the cytoplasm (I_cyt_).

For NLS intensity analysis (**Figure S9**), rectangular intensity profiles (1 μm width) were taken along the cell length. Fluorescent intensity was normalized with respect to peak intensity of the corresponding profile and centered with respect to the nucleus midpoint.

### 5.11 Calculation of cell adhesion forces

Using our previously reported Nanonet Force Microscopy technique, cell-fiber adhesion forces were quantified using a custom MATLAB code. Briefly, taut nanofibers were approximated as loaded Euler Bernoulli beams with fixed-fixed boundary conditions and with forces exerted at either cell pole (localization of the focal adhesion clusters) at an angle. This direction of force exertion was taken along the average orientation (with respect to the undeflected fibers) of the f-actin stress fibers emanating from the polar FA clusters. Next, experimentally observed nanofiber deflection profiles were compared with the best fit profile from finite-element model predictions and the error between them was minimized iteratively to generate the adhesion forces. In fiber doublets (homogenous diameters), due to the high level of symmetry in fiber deflections on either fiber, magnitude of the net cell force exerted is approximated as twice the magnitude of the cell-adhesion force with each fiber.

### 5.12 Statistical analysis and data fitting

Statistical analysis was performed in GraphPad Prism (GraphPad Software, La Jolla, CA, USA) software. For statistical comparison between multiple groups, ANOVA (along with Tukey’s honestly significant difference test) was utilized. For pairwise comparisons with a control group, Student’s t-test was used. In all scatter data column plots, error bars represent standard deviation. Unless otherwise mentioned, in all other plots, error bars represent standard error of measurement. For all statistical plots *,**,***,**** represent p< 0.05, 0.01, 0.001 and 0.0001 respectively. Data fitting was performed in either MATLAB (for nonlinear fits) or Microsoft Excel (linear fits).

## Supporting Information

Supplemental Figures S1–S11. Movies 1-11.
Supporting Information is available from the Wiley Online Library or from the author.
Supplemental Figures S1–S11.
Movie M1 (.MP4): C2C12 (mouse myoblasts) protruding on fiber doublets during spreading (timestamp-h:min:sec).
Movie M2 (.MP4): C2C12 (mouse myoblasts) spreading on 200 nm diameter fiber doublets
Movie M3 (.MP4): C2C12 (mouse myoblasts) spreading on 350 nm diameter fiber doublets
Movie M4 (.MP4): C2C12 (mouse myoblasts) spreading on 800 nm diameter fiber doublets
Movie M5 (.MP4): C2C12 spreading on 200 nm-800 nm mismatch diameter fiber doublets
Movie M6 (.MP4): C2C12 spreading on 200 nm-800 nm-200 nm mismatch diameter fiber triplet
Movie M7 (.MP4): C2C12 spreading on 800 nm-200 nm-800 nm mismatch diameter fiber triplet
Movie M8 (.MP4): C2C12 spreading on single 350 nm diameter fibers (timestamp-h:min:sec).
Movie M9 (.MP4): C2C12 cells expressing NLS mCherry spreading on single 200 nm diameter fiber
Movie M10 (.MP4): C2C12 spreading on three 350 nm diameter fibers.
Movie M11 (.MP4): Two C2C12 cells transition from 4-fibers to 7-fibers (350 nm diameter) during spreading.
All timestamps are in hours : minutes format unless otherwise specified.

## Acknowledgements

The authors thank members of the Spinneret-based Tunable Engineered Parameters (STEP) Lab, Virginia Tech and Konstantopoulos Lab, Johns Hopkins University for helpful suggestions and discussions. HeLa cells were kindly provided by Dr. Jennifer DeLuca, Colorado State University. The authors thank the Institute of Critical Technologies and Sciences (ICTAS) and Macromolecules Innovation Institute at Virginia Tech for the support to conduct this study. ASN acknowledges partial support for this study from National Science Foundation (NSF grant # 1762468) and KK acknowledges support from National Institute of Health (NIH grants CA183804 and RO1 CA254193).

## Author contributions

AG and AT are equally contributing second authors. ASN conceived and oversaw the project. ASN and KK designed experiments. AJ, AT, and AG performed experiments. AJ, AT, AG, KK, and ASN analyzed data. AJ and AT wrote the manuscript. RKK and ASN developed the cell force model. AT and KK developed the lamin deficient cell lines. AJ and AG developed the suspended fiber networks. AJ performed confocal imaging. All authors contributed to reading and editing the manuscript.

## Supporting Information

**Figure S1.**
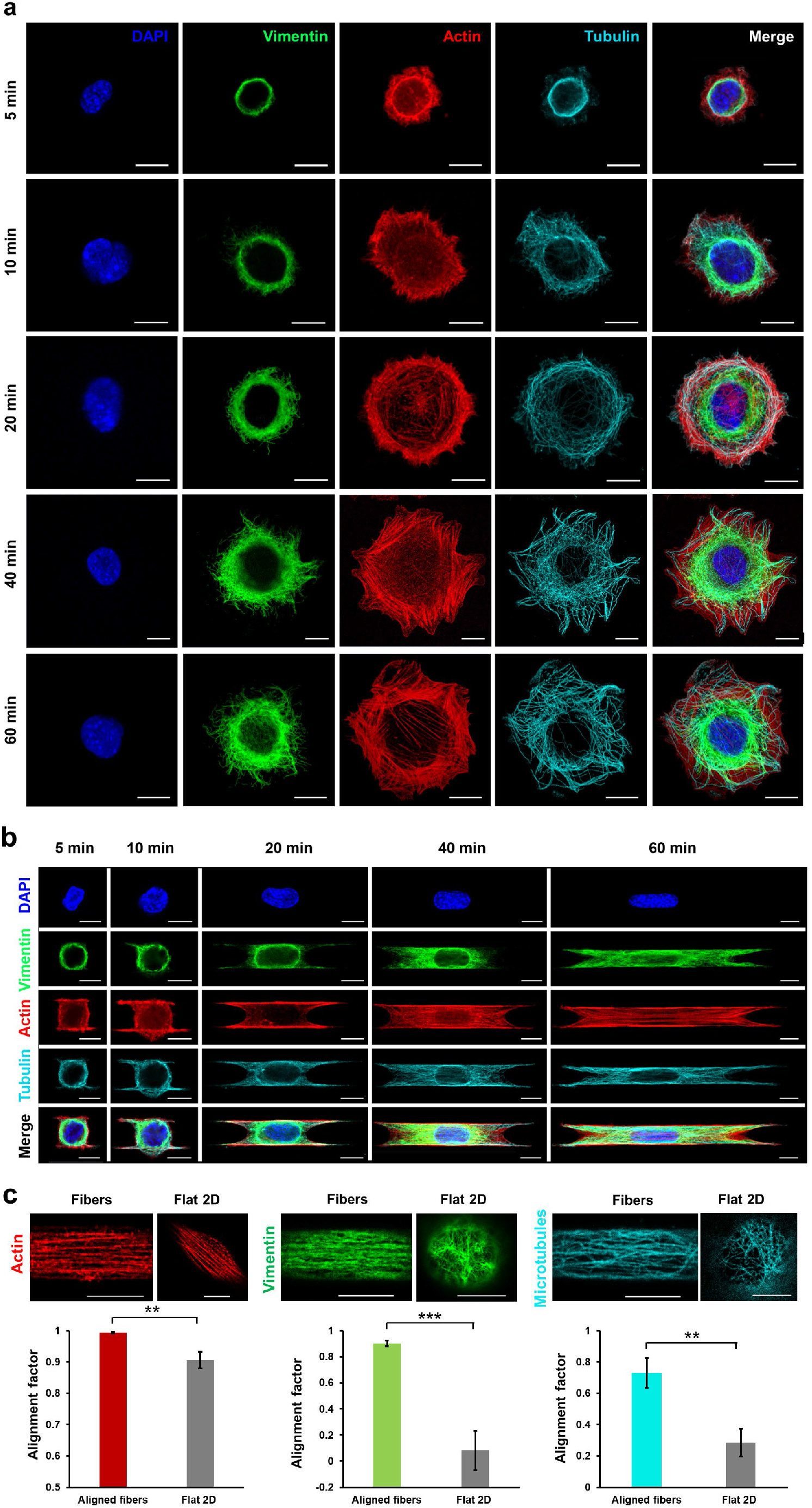
Cytoskeletal alignment in suspended fibers. a,b) Representative cells showing cytoskeletal organization at different timepoints during spreading on flat glass and suspended fiber doublets (fiber diameter: 350 nm) respectively, cells are stained for actin (red), vimentin intermediate filaments (green), microtubules (cyan) and nuclei (blue), cells on fibers show entry of microtubules and intermediate filaments into the protrusions as spreading progresses, scale bars represent 10 μm c) Comparison of the alignment of the cytoskeletal elements in the perinuclear region, between flat glass and suspended fibers (fiber diameter: 350 nm). Alignment factor is defined as cos(2α) where α is the orientation of the actin stress fiber or individual pores within the microtubule or intermediate filament meshwork, with respect to the primary cell orientation, Alignment factor ~ 1 for perfect alignment and ~ 0 for random alignment, scale bars represent 10 μm, n=5-6 cells for aligned fibers and Flat 2D respectively.

**Figure S2.**
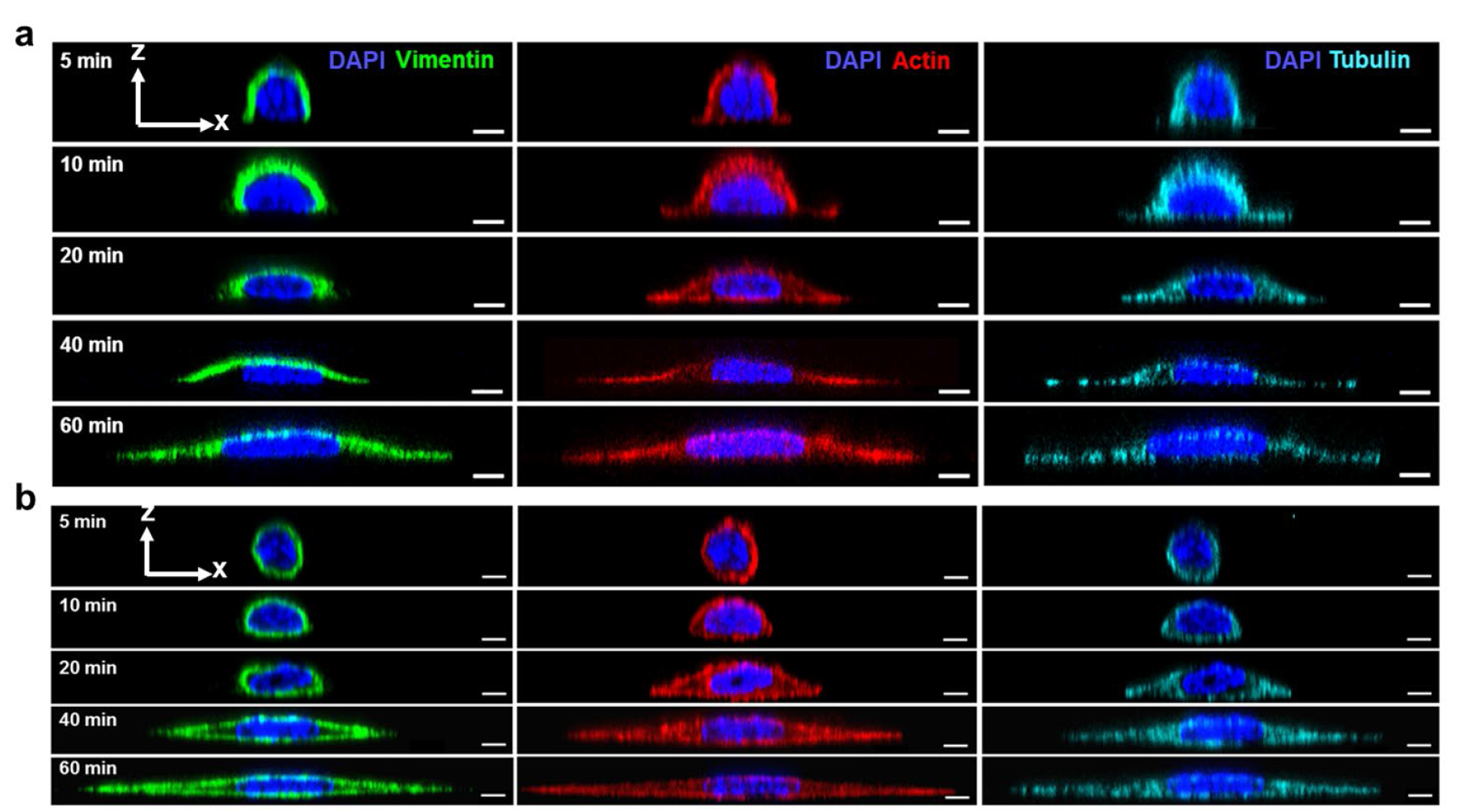
Spatiotemporal localization of cytoskeletons. a) Confocal side views (xz) showing ‘Capping’ localization of the cytoskeletal elements at various timepoints during cell spreading on flat glass b) ‘Caging’ localization observed in suspended fibers (fiber diameter: 350 nm), actin (red), vimentin intermediate filaments (green), microtubules (cyan) and nuclei (blue), all scale bars represent 5 μm

**Figure S3.**
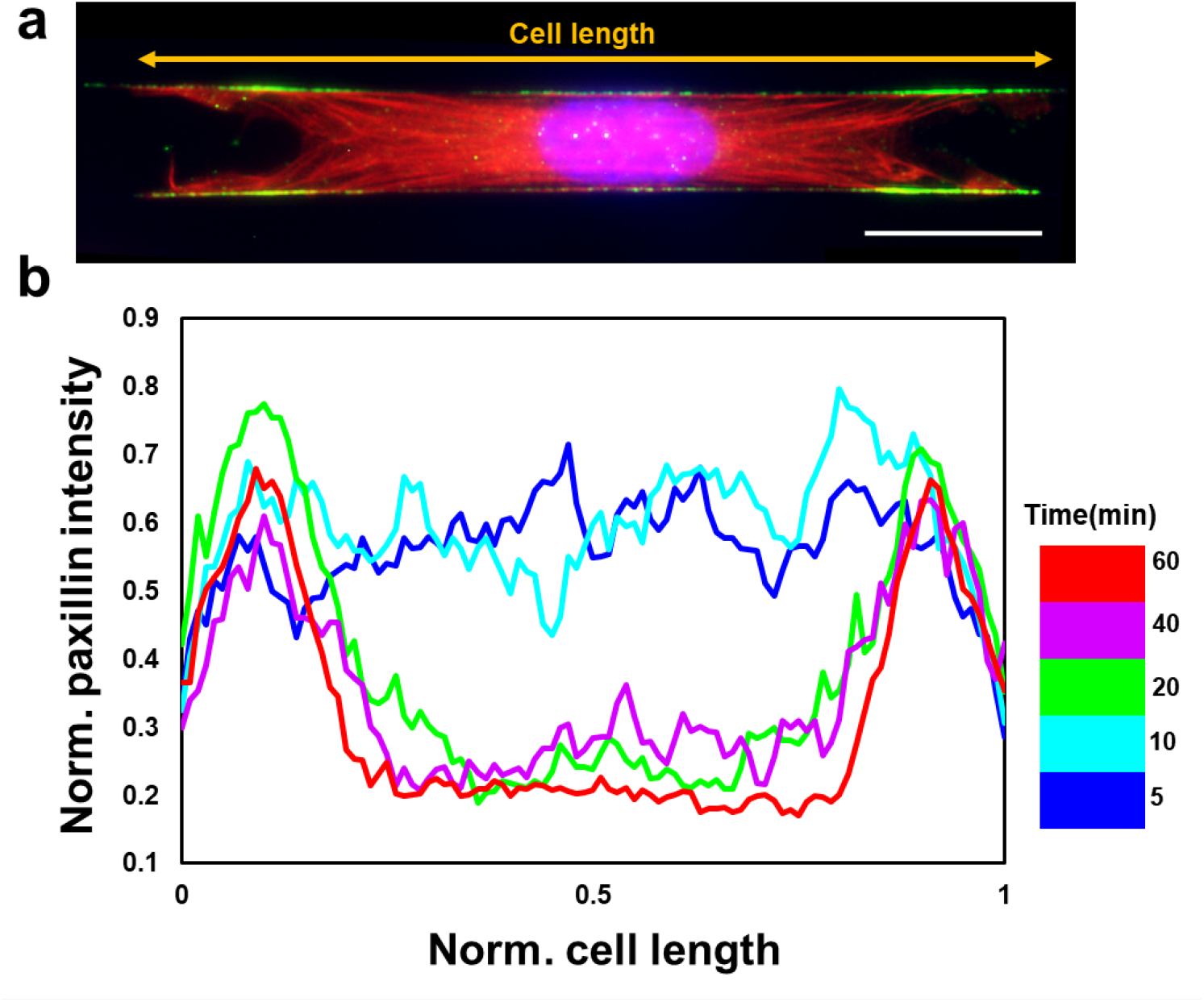
Evolution of focal adhesion organization over the course of cell spreading. a) Representative spread cell showing focal adhesion clustering at cell poles, scale bars represent 20 μm, b), Average profiles (n=9-12 for each timepoint) are shown, Paxillin intensity is normalized with respect to the peak intensity of each profile, Cell length normalization was performed with respect to the entire length of the cell as shown in part a, In the spread state, two intensity peaks demonstrate the focal adhesion clustering, Data is shown for 350 nm diameter category

**Figure S4.**
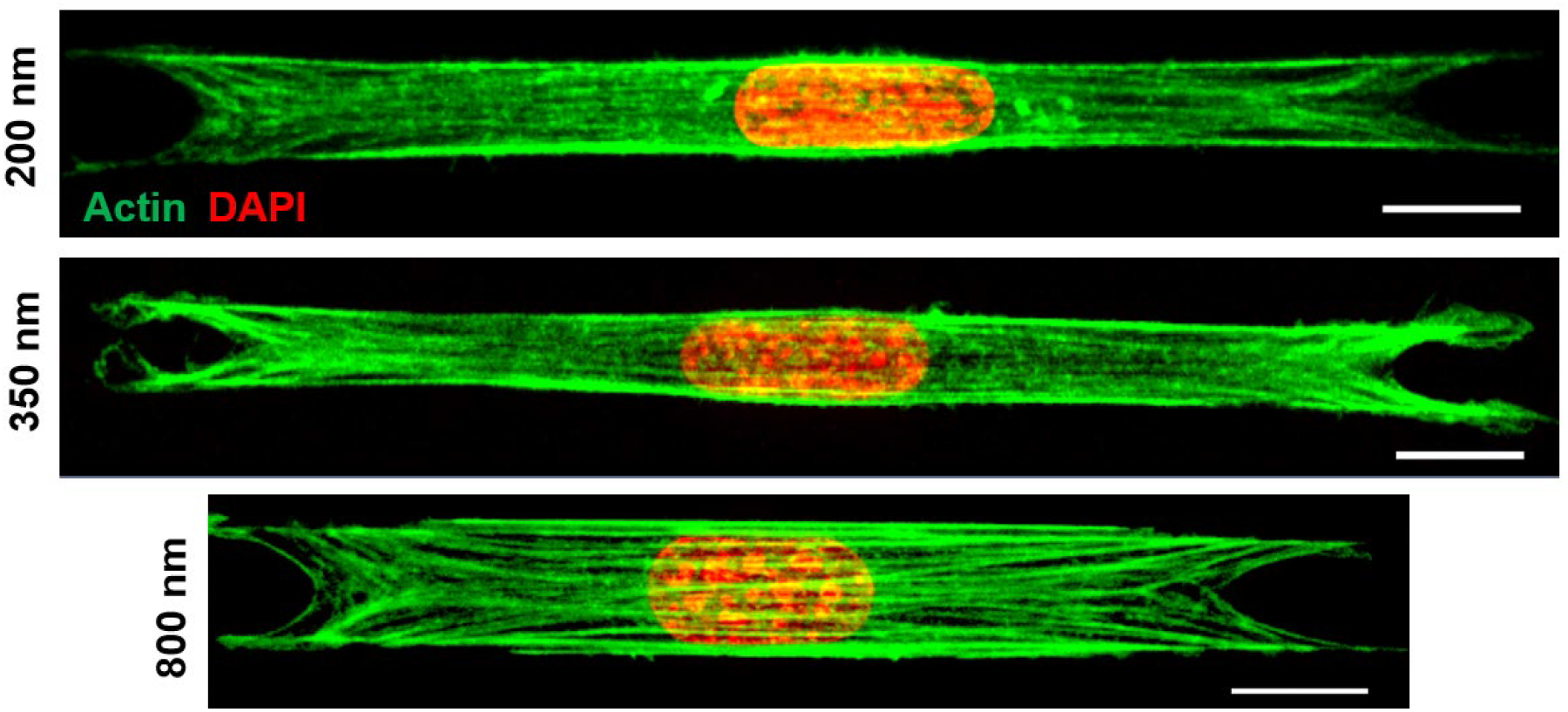
Stress fiber organization in spread cells, Maximum intensity projections of representative cells demonstrating stress fibers emerge from focal adhesion clusters on either side and converge to become almost parallel to the cell orientation, in the perinuclear region. Actin and DAPI are pseudo-colored in green and red for better visualization. Scale bars represent 10 μm.

**Figure S5.**
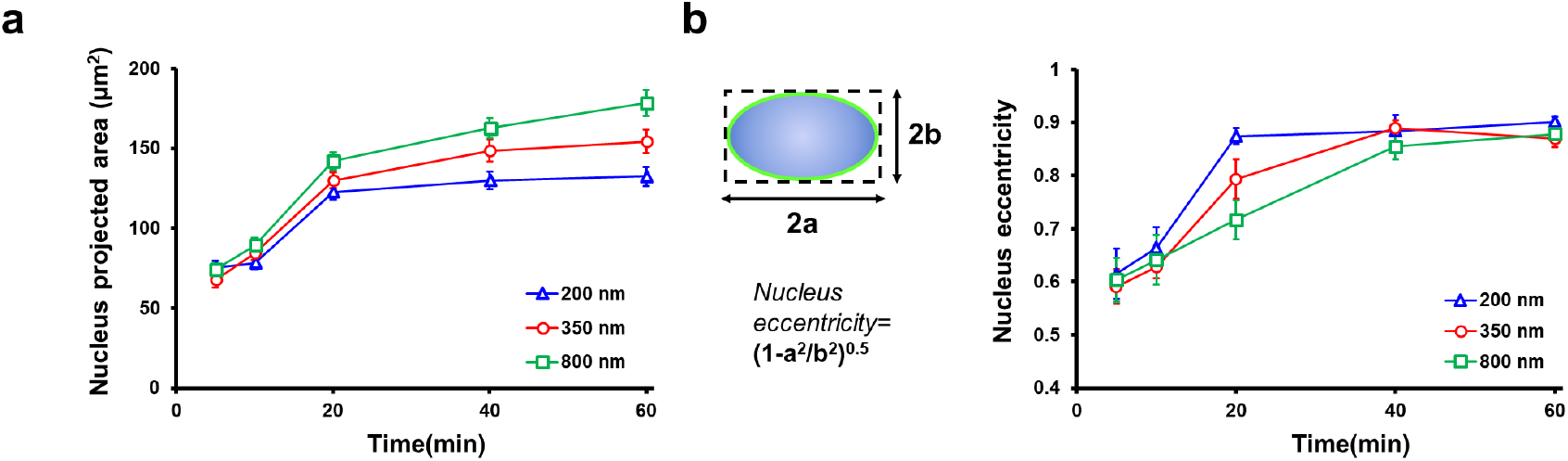
Nucleus geometry during cell spreading. a,b) Evolution of nucleus projected area (xy) and eccentricity as function of spreading time and shown for the different diameter fiber doublets, respectively, n=17-34 per timepoint for each diameter category

**Figure S6.**
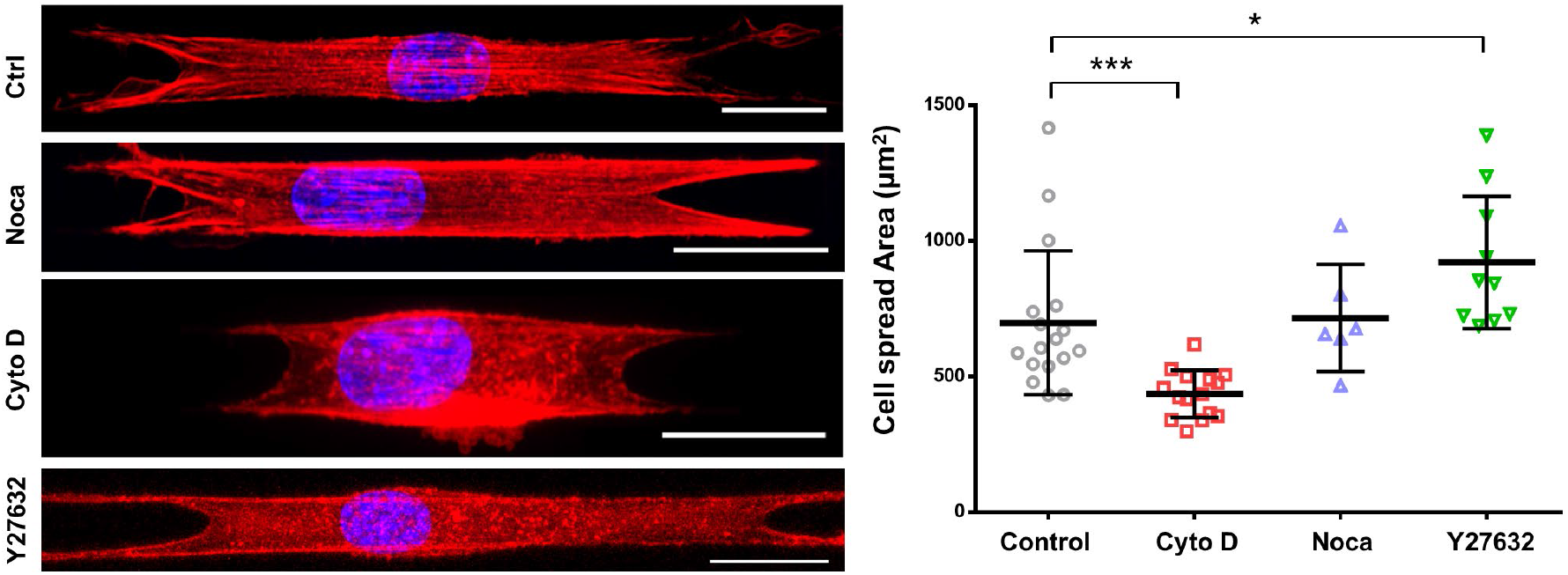
Effect of cytoskeletal inhibition on cell spreading, Representative images showing control (no treatment), cytochalasin D, nocodazole and Y27632 treated cells, fiber diameter: 350 nm, scale bars represent 20 μm. Comparison of the cell spread area under different drug conditions, n=17,15,6 and 10.

**Figure S7.**
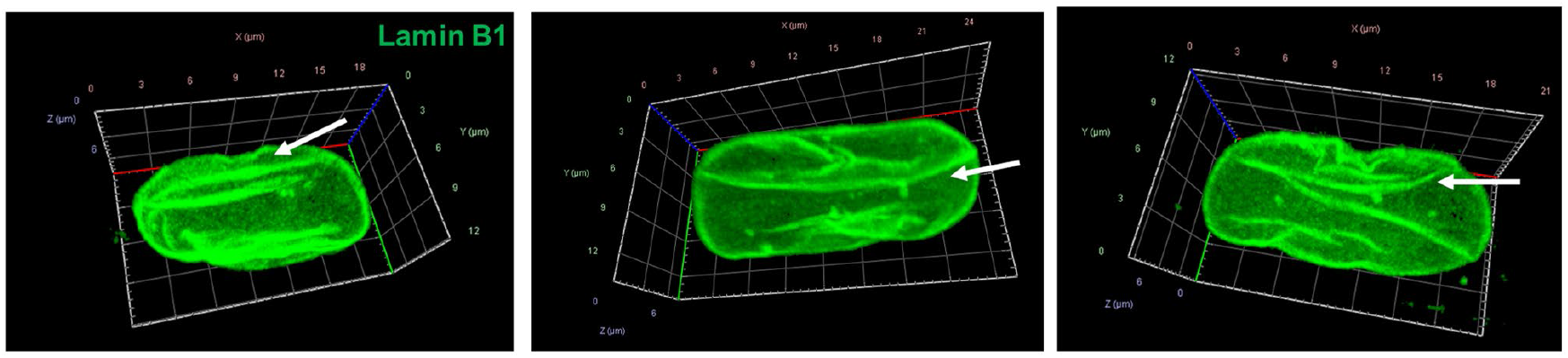
Morphological alterations in Lamin A/C KD cells. a) 3D isometric views showing wrinkled morphologies of Lamin A/C KD cells, white arrows indicate wrinkles, Lamin B1 is shown in green

**Figure S8.**
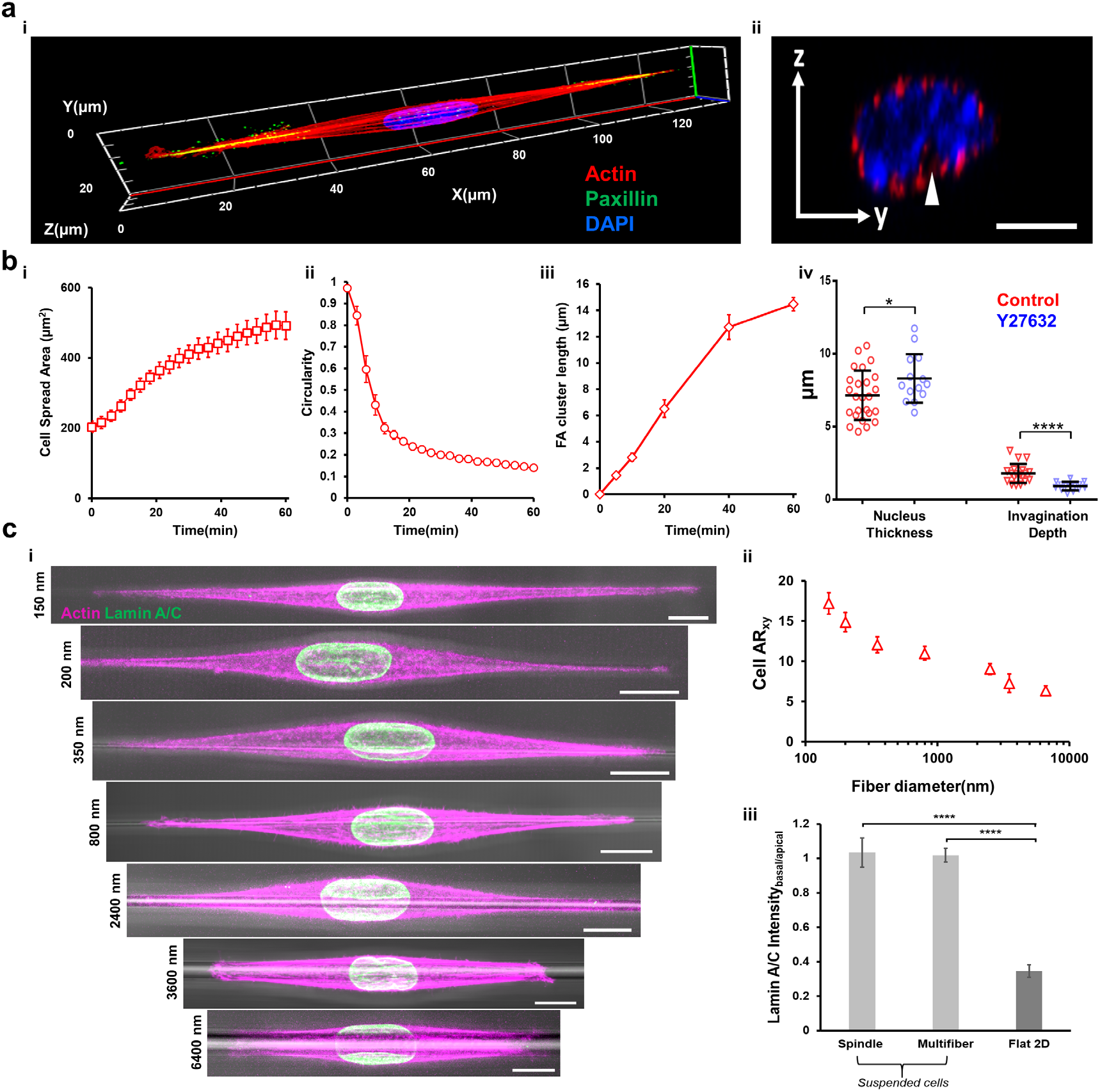
Cell and nucleus shapes in spindle cells: a) i) 3D isometric view of spindle cell stained for actin (red), paxillin (green, shown with yellow arrows) and nucleus (blue) ii) Representative confocal side views (yz) showing stress-fiber mediated caging of nucleus in spindle shaped cells, scale bars represent 5 μm b) i.ii) Temporal evolution of cell spread area and circularity for spindle cells, n=15 cells, fiber diameter: 350 nm, iii) Focal adhesion cluster (FAC) length evolution during cell spreading, n=34.47,32,17 and 34 for 5, 10,20,40 and 60 min respectively, iv) Effect of cell contractility inhibition on nucleus thickness and invagination size in spindle cells, (fiber diameter: 350 nm) c) i) Representative images showing variation of cell/nucleus shape as the fiber diameter is varied from 150-6600 nm. Actin (magenta) and lamin A/C (green) are overlaid with the brightfield channel to show underlying fibers, scale bars represent 10 μm ii) Cell aspect ratio as a function of fiber diameter, n=14,10,10,11,10,6 and 6 for 150,200,350,800,2500,3500 and 6600 nm respectively iii) Localization of lamin A/C at the basal (contact points with fiber) and apical surfaces for spindle (1-fiber) and multifiber configurations, Flat 2D is included for comparisons (n=5-10 cells).

**Figure S9.**
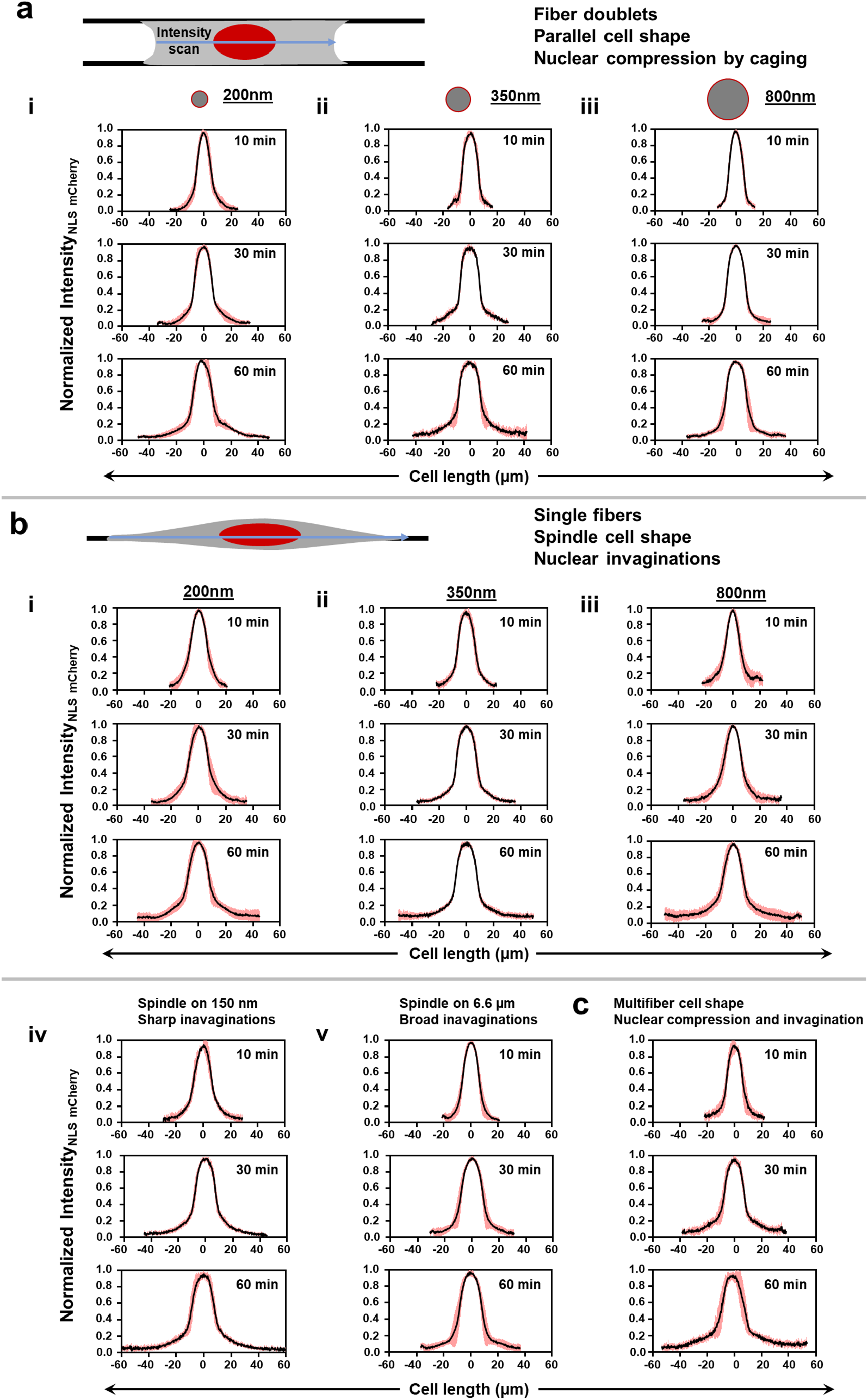
NLS localization in aligned fibers. a) i,ii,iii) NLS intensity profiles, along cell length at different spreading timepoints (~ 10,30,60 min) corresponding to fiber doublets with 200 nm, 350 nm and 800 nm respectively, Schematic illustrates the direction of intensity scans b) i,ii,iii, iv and v) Spatiotemporal NLS localization for spindle shaped cells corresponding to fiber diameters: 200, 350,800, 150 and 6600 nm respectively c) Intensity analysis for multifiber (D: 350 nm) cells, n=10 cells for each timepoint and substrate category. Error bars represent standard deviation.

**Figure S10.**
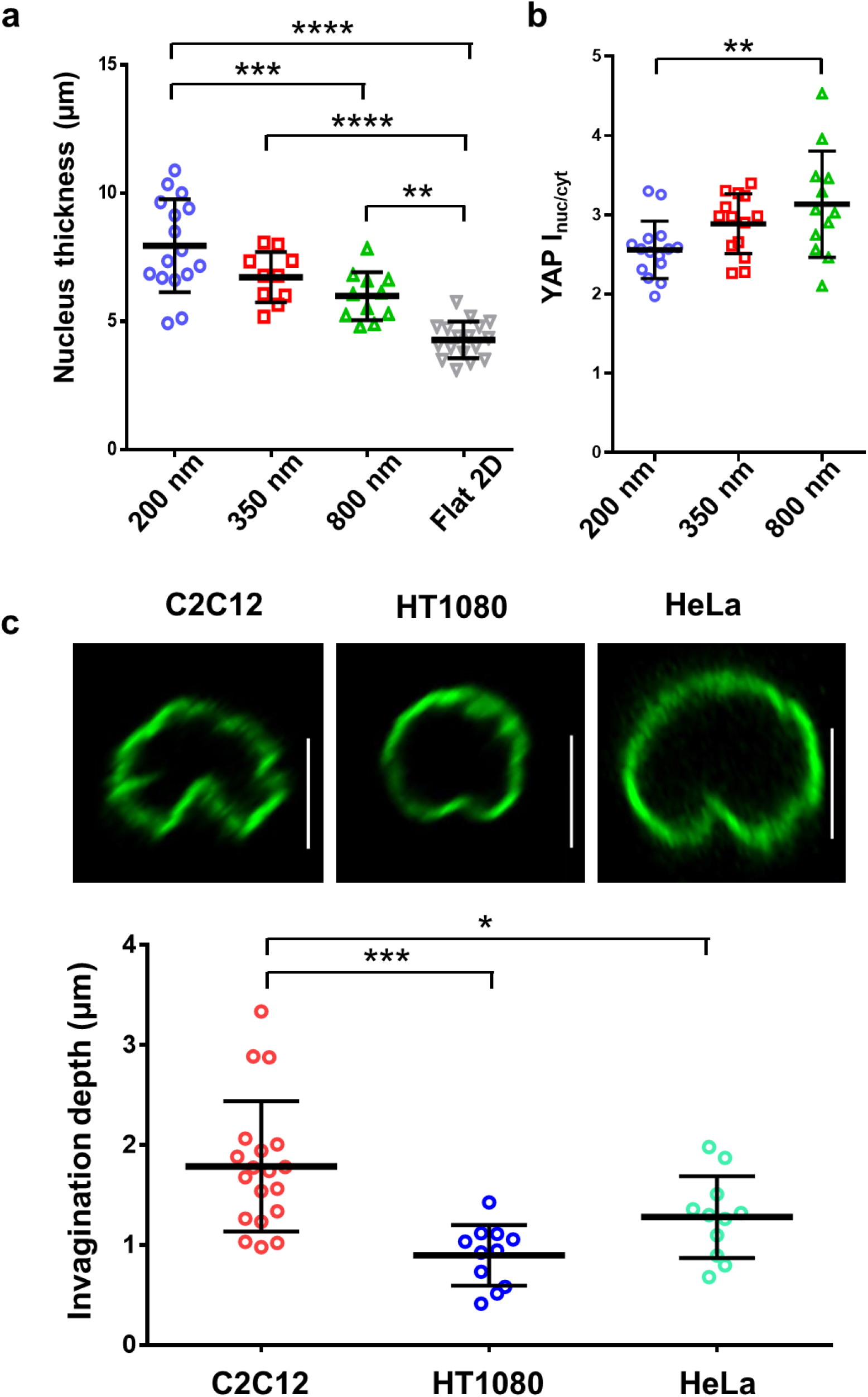
Nucleus shape regulation mode is independent of cell line: a) Fiber diameter dependence of nucleus thickness in HT1080 fibrosarcoma cells b) Enhanced nuclear translocation of YAP for HT1080 cells in 800 nm networks. c) Nuclear invaginations are observed in spindle cells for all cell types tested (C2C12, HT1080 and HeLa), comparison of the invagination depths in the different cell types for 350 nm diameter single fibers, scale bars represent 5 μm.

**Figure S11:**
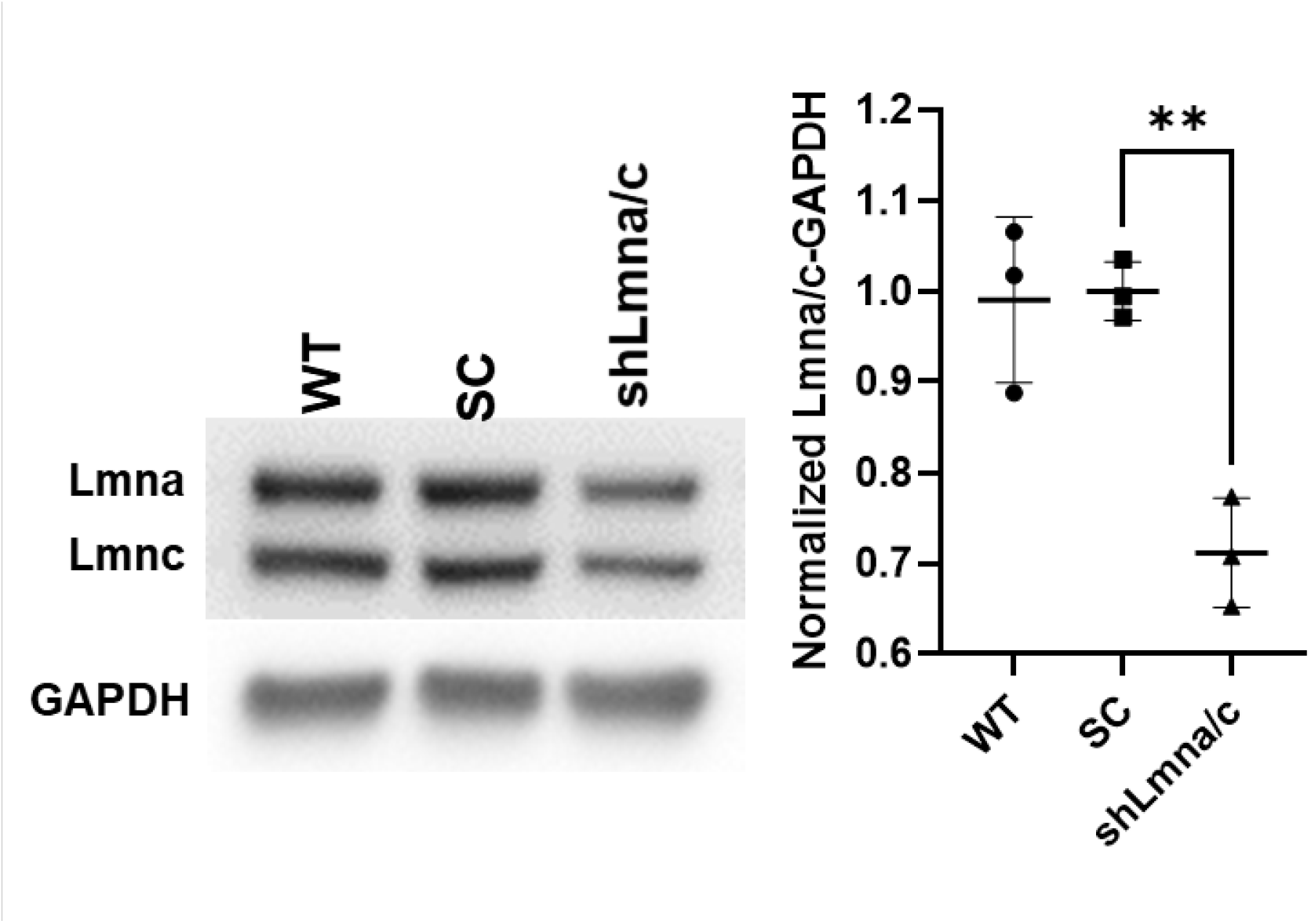
Western blot showing partial knockdown of Lamin A/C in C2C12 cells. SC stands for scramble with null expression.

## References

[1] F. Guilak, J. R. Tedrow, R. Burgkart, Biochem. Biophys. Res. Commun. 2000, 269, 781.

[2] J. Swift, D. E. Discher, J. Cell Sci. 2014, 127, 3005.

[3] B. M. Skinner, E. E. P. Johnson, Chromosoma 2017, 126, 195.

[4] C. Uhler, G. V. Shivashankar, Trends in Cancer 2018, 4, 320.

[5] S. G. Alam, D. Lovett, D. I. Kim, K. J. Roux, R. B. Dickinson, T. P. Lele, J. Cell Sci. 2015, 128, 1901.

[6] M. L. Lombardi, D. E. Jaalouk, C. M. Shanahan, B. Burke, K. J. Roux, J. Lammerding, J. Biol. Chem. 2011, 286, 26743.

[7] T. P. Lele, R. B. Dickinson, G. G. Gundersen, J. Cell Biol. 2018, 217, 3330.

[8] A. D. Stephens, P. Z. Liu, E. J. Banigan, L. M. Almassalha, V. Backman, S. A. Adam, R. D. Goldman, J. F. Marko, Mol. Biol. Cell 2018, 29, 220.

[9] Y. Xia, I. L. Ivanovska, K. Zhu, L. Smith, J. Irianto, C. R. Pfeifer, C. M. Alvey, J. Ji, D. Liu, S. Cho, R. R. Bennett, A. J. Liu, R. A. Greenberg, D. E. Discher, J. Cell Biol. 2018, 217, 3796.

[10] W. H. De vos, F. Houben, M. Kamps, A. Malhas, F. Verheyen, J. Cox, E. M. M. Manders, V. L. R. M. Verstraeten, M. A. M. Van steensel, C. L. M. Marcelis, A. Van den wijngaard, D. J. Vaux, F. C. S. Ramaekers, J. L. V. Broers, Hum. Mol. Genet. 2011, 20, 4175.

[11] F. Alisafaei, D. S. Jokhun, G. V. Shivashankar, V. B. Shenoy, Proc. Natl. Acad. Sci. 2019, 116, 13200.

[12] A. Elosegui-Artola, I. Andreu, A. E. M. Beedle, A. Lezamiz, M. Uroz, A. J. Kosmalska, R. Oria, J. Z. Kechagia, P. Rico-Lastres, A. L. Le Roux, C. M. Shanahan, X. Trepat, D. Navajas, S. Garcia-Manyes, P. Roca-Cusachs, Cell 2017, 171, 1397.

[13] T. J. Kirby, J. Lammerding, Nat. Cell Biol. 2018, 20, 373.

[14] K. Damodaran, S. Venkatachalapathy, F. Alisafaei, A. V. Radhakrishnan, D. S. Jokhun, V. B. Shenoy, G. V. Shivashankar, Mol. Biol. Cell 2018, 29, 3039.

[15] P. Mistriotis, E. O. Wisniewski, K. Bera, J. Keys, Y. Li, S. Tuntithavornwat, R. A. Law, N. A. Perez-Gonzalez, E. Erdogmus, Y. Zhang, R. Zhao, S. X. Sun, P. Kalab, J. Lammerding, K. Konstantopoulos, J. Cell Biol. 2019, 218, 4093.

[16] B. R. Freedman, A. B. Rodriguez, R. J. Leiphart, J. B. Newton, E. Ban, J. J. Sarver, R. L. Mauck, V. B. Shenoy, L. J. Soslowsky, Sci. Rep. 2018, 8, 1.

[17] P. P. Provenzano, R. Vanderby, Matrix Biol. 2006, 25, 71.

[18] T. A. H. Järvinen, L. Józsa, P. Kannus, T. L. N. Järvinen, M. Järvinen, J. Muscle Res. Cell Motil. 2002, 23, 245.

[19] A. R. Gillies, R. L. Lieber, Muscle and Nerve 2011, 44, 318.

[20] M. W. Conklin, J. C. Eickhoff, K. M. Riching, C. A. Pehlke, K. W. Eliceiri, P. P. Provenzano, A. Friedl, P. J. Keely, Am. J. Pathol. 2011, 178, 1221.

[21] P. P. Provenzano, D. R. Inman, K. W. Eliceiri, J. G. Knittel, L. Yan, C. T. Rueden, J. G. White, P. J. Keely, BMC Med. 2008, 6, 11.

[22] K. R. Levental, H. Yu, L. Kass, J. N. Lakins, M. Egeblad, J. T. Erler, S. F. T. Fong, K. Csiszar, A. Giaccia, W. Weninger, M. Yamauchi, D. L. Gasser, V. M. Weaver, Cell 2009, 139, 891.

[23] J. M. Szulczewski, D. R. Inman, M. Proestaki, J. Notbohm, B. M. Burkel, S. M. Ponik, Acta Biomater. 2021, DOI 10.1016/j.actbio.2021.04.053.

[24] J. Starke, K. Maaser, B. Wehrle-Haller, P. Friedl, Exp. Cell Res. 2013, 319, 2424.

[25] A. D. Doyle, N. Carvajal, A. Jin, K. Matsumoto, K. M. Yamada, Nat. Commun. 2015, 6, DOI 10.1038/ncomms9720.

[26] T. Ushiki, Arch. Histol. Cytol. 2002, 65, 109.

[27] M. Fernández, J. Keyriläinen, R. Serimaa, M. Torkkeli, M. L. Karjalainen-Lindsberg, M. Tenhunen, W. Thomlinson, V. Urban, P. Suortti, Phys. Med. Biol. 2002, 47, 577.

[28] A. Padhi, A. S. Nain, Ann. Biomed. Eng. 2020, 48, DOI 10.1007/s10439-019-02337-7.

[29] P. Keely, A. Nain, F1000Research 2015, 4, DOI 10.12688/f1000research.6623.1.

[30] K. Wolf, M. te Lindert, M. Krause, S. Alexander, J. te Riet, A. L. Willis, R. M. Hoffman, C. G. Figdor, S. J. Weiss, P. Friedl, J. Cell Biol. 2013, 201, 1069.

[31] P. Friedl, E. Sahai, S. Weiss, K. M. Yamada, Nat. Rev. Mol. Cell Biol. 2012, 13, 743.

[32] A. D. Doyle, F. W. Wang, K. Matsumoto, K. M. Yamada, J. Cell Biol. 2009, 184, 481.

[33] S. Meehan, A. S. Nain, Biophys. J. 2014, 107, 2604.

[34] B. Koons, P. Sharma, Z. Ye, A. Mukherjee, M. H. Lee, D. Wirtz, B. Behkam, A. S. Nain, ACS Nano 2017, 11, 12037.

[35] A. Mukherjee, B. Behkam, A. S. Nain, iScience 2019, 19, 905.

[36] J. Singh, A. Pagulayan, B. A. Camley, A. S. Nain, Proc. Natl. Acad. Sci. 2021, 118, e2011815118.

[37] A. Jana, I. Nookaew, J. Singh, B. Behkam, A. T. Franco, A. S. Nain, FASEB J. 2019, 33, 10618.

[38] K. Sheets, J. Wang, W. Zhao, R. Kapania, A. S. Nain, Biophys. J. 2016, 111, 197.

[39] K. Sheets, S. Wunsch, C. Ng, A. S. Nain, Acta Biomater. 2013, 9, 7169.

[40] B. Tu-Sekine, A. Padhi, S. Jin, S. Kalyan, K. Singh, M. Apperson, R. Kapania, S. C. Hur, A. Nain, S. F. Kim, FASEB J. 2019, 33, 14137.

[41] P. M. Graybill, A. Jana, R. K. Kapania, A. S. Nain, R. V. Davalos, ACS Nano 2021, 15, 2554.

[42] A. Padhi, K. Singh, J. Franco-Barraza, D. J. Marston, E. Cukierman, K. M. Hahn, R. K. Kapania, A. S. Nain, Commun. Biol. 2020, 3, 390.

[43] H. Wolfenson, T. Iskratsch, M. P. Sheetz, Biophys. J. 2014, 107, 2508.

[44] Y. Li, D. Lovett, Q. Zhang, S. Neelam, R. A. Kuchibhotla, R. Zhu, G. G. Gundersen, T. P. Lele, R. B. Dickinson, Biophys. J. 2015, 109, 670.

[45] A. Katiyar, V. J. Tocco, Y. Li, V. Aggarwal, A. C. Tamashunas, R. B. Dickinson, T. P. Lele, Soft Matter 2019, 15, 9310.

[46] S. B. Khatau, C. M. Hale, P. J. Stewart-Hutchinson, M. S. Patel, C. L. Stewart, P. C. Searson, D. Hodzic, D. Wirtz, Proc. Natl. Acad. Sci. U. S. A. 2009, 106, 19017.

[47] T. O. Ihalainen, L. Aires, F. A. Herzog, R. Schwartlander, J. Moeller, V. Vogel, Nat. Mater. 2015, 14, 1252.

[48] S. Dupont, L. Morsut, M. Aragona, E. Enzo, S. Giulitti, M. Cordenonsi, F. Zanconato, J. Le Digabel, M. Forcato, S. Bicciato, N. Elvassore, S. Piccolo, Nature 2011, 474, 179.

[49] J. K. Kim, A. Louhghalam, G. Lee, B. W. Schafer, D. Wirtz, D. H. Kim, Nat. Commun. 2017, 8, 1.

[50] A. D. Stephens, E. J. Banigan, J. F. Marko, Curr. Opin. Cell Biol. 2019, 58, 76.

[51] A. S. Nain, M. Sitti, A. Jacobson, T. Kowalewski, C. Amon, Macromol. Rapid Commun. 2009, 30, 1406.

[52] A. S. Nain, J. Wang, Polym. J. 2013, 45, 695.

[53] J. Wang, A. S. Nain, Langmuir 2014, 30, DOI 10.1021/la503011u.

[54] A. Buxboim, J. Irianto, J. Swift, A. Athirasala, J. W. Shin, F. Rehfeldt, D. E. Discher, Mol. Biol. Cell 2017, 28, 3333.

[55] P. G. Gritsenko, O. Ilina, P. Friedl, J. Pathol. 2012, 226, 185.

[56] P. Friedl, K. Wolf, J. Cell Biol. 2010, 188, 11.

[57] A. D. Doyle, F. W. Wang, K. Matsumoto, K. M. Yamada, J. Cell Biol. 2009, 184, 481.

[58] S. P. Carey, C. M. Kraning-Rush, R. M. Williams, C. a Reinhart-King, Biomaterials 2012, 33, 4157.

[59] C. A. Reinhart-King, M. Dembo, D. A. Hammer, Biophys. J. 2005, 89, 676.

[60] M. Théry, A. Pépin, E. Dressaire, Y. Chen, M. Bornens, Cell Motil. Cytoskeleton 2006, 63, 341.

[61] E. Cukierman, Science (80-. ). 2001, 294, 1708.

[62] K. M. Hakkinen, J. S. Harunaga, A. D. Doyle, K. M. Yamada, Tissue Eng. Part A 2011, 17, 713.

[63] N. Q. Balaban, U. S. Schwarz, D. Riveline, P. Goichberg, G. Tzur, I. Sabanay, D. Mahalu, S. Safran, A. Bershadsky, L. Addadi, B. Geiger, Nat. Cell Biol. 2001, 3, 466.

[64] M. Versaevel, T. Grevesse, S. Gabriele, Nat. Commun. 2012, 3, 671.

[65] S. Sugita, T. Adachi, Y. Ueki, M. Sato, Biophys. J. 2011, 101, 53.

[66] K. Katoh, Y. Kano, M. Masuda, H. Onishi, K. Fujiwara, Mol. Biol. Cell 1998, 9, 1919.

[67] D. B. Lovett, N. Shekhar, J. a. Nickerson, K. J. Roux, T. P. Lele, Cell. Mol. Bioeng. 2013, DOI 10.1007/s12195-013-0270-2.

[68] B. Sen, G. Uzer, R. M. Samsonraj, Z. Xie, C. McGrath, M. Styner, A. Dudakovic, A. J. van Wijnen, J. Rubin, Stem Cells 2017, 35, 1624.

[69] P. M. Davidson, H. Özçelik, V. Hasirci, G. Reiter, K. Anselme, Adv. Mater. 2009, 21, 3586.

[70] F. Badique, D. R. Stamov, P. M. Davidson, M. Veuillet, G. Reiter, J. N. Freund, C. M. Franz, K. Anselme, Biomaterials 2013, 34, 2991.

[71] C. M. Denais, R. M. Gilbert, P. Isermann, A. L. McGregor, M. Te Lindert, B. Weigelin, P. M. Davidson, P. Friedl, K. Wolf, J. Lammerding, Science (80-. ). 2016, 352, 353.

